# Analysis and control of untemplated DNA polymerase activity for guided synthesis of kilobase-scale DNA sequences

**DOI:** 10.1101/2024.08.29.610422

**Authors:** Simeon D. Castle, Adrian Woolfson, Gregory Linshiz, Blake T. Riley, Ifor D.W. Samuel, Philipp Holliger, Lauren Oldfield, Andrew Hessel, Thomas E. Gorochowski

## Abstract

DNA polymerases are complex molecular machines able to replicate genetic material using a template-driven process. While the copying function of these enzymes is well established, their ability to perform untemplated DNA synthesis is less well characterized. Here, we explore the ability of DNA polymerases to synthesize DNA fragments in the absence of template. We use long-read nanopore sequencing and real-time PCR to observe the synthesis of pools of DNA products derived from a diverse set of natural and engineered DNA polymerases across varying temperatures and buffer compositions. We detail the features of the DNA fragments generated, enrichment of select sequence motifs, and demonstrate that the sequence composition of the synthesized DNA may be altered by modifying environmental conditions. This work provides an extensive data set to better discern the process of untemplated DNA polymerase activity and may support its potential repurposing as a technology for the guided synthesis of DNA sequences on the kilobase-scale and beyond.

## Introduction

Genetic information encoded in DNA sequences is a defining feature of all life on Earth. It provides the instructions that cells require to synthesise and assemble their molecular machinery, encodes signals to coordinate cellular processes, and is the substrate for processes of evolution by natural selection. How new biological information emerges however, remains uncertain.

DNA polymerases (DNAPs) are typically viewed as molecular machines involved in the high fidelity copying of genetic information through the replication of DNA using a template-mediated process ^1^. However, it was shown in the 1960s that some DNAPs have the capacity to synthesise single-stranded DNA fragments in the absence of a pre-existing template ^2–4^. This *ab initio* or “doodling” activity is widespread among polymerases ^5^ and has been observed in RNA polymerases, notably T7 RNA polymerase ^6^, Qbeta RNA-dependent RNA polymerase^7^ and 5TU RNA polymerase ribozyme ^8^. Doodling suggests a mechanism for how *de novo* genetic information may be generated.

Investigations of this process have demonstrated that DNAPs have an intrinsic capacity to doodle^9–12^, that this process is sensitive to environmental factors such as temperature and variations in buffer conditions ^2,9,10,13^, and that pools of synthesised DNA molecules produced in the absence of template likely go through a multi-stage process of semi-random synthesis and enrichment via the selection of sequences able to more efficiently self-replicate ^5,14–16^. However, all such studies have been hampered by an inability to extensively characterise the pools of DNA molecules generated. Recent advances in the ability to read full-length DNA molecules using long-read sequencing have made it feasible to generate near-complete sequence information from such vast pools, thereby gaining new insight into the generative process.

In this work, we employ nanopore sequencing and real-time PCR to provide the most comprehensive characterisation to date of both doodling kinetics and sequence features of the full-length untemplated DNA molecules synthesized (**Figure 1a**). We document the doodling function of a diverse array of natural and engineered DNAPs, assess how temperature and buffer composition impacts the nature of the sequences generated, and explore how this process may potentially be manipulated through limiting deoxynucleotide triphosphate (dNTP) availability. We identify considerably greater diversity in the features of the sequences generated than previously reported, document the complexity of the DNA molecules synthesised, and demonstrate that in select cases, modification of environmental conditions enables robust and directed changes to the sequences produced. In addition to providing a rich data set for further characterizing the nature of untemplated DNA synthesis, our findings also suggest a need to reconsider the role of DNAP function in cellular biochemistry. Overall, the work offers a new perspective on how DNAPs may contribute to the genesis and evolution of new genetic information and may provide the basis of a method for repurposing DNAPs as a platform for programming the synthesis of DNA sequences on the kilobase-scale and beyond.

**Figure 1:**
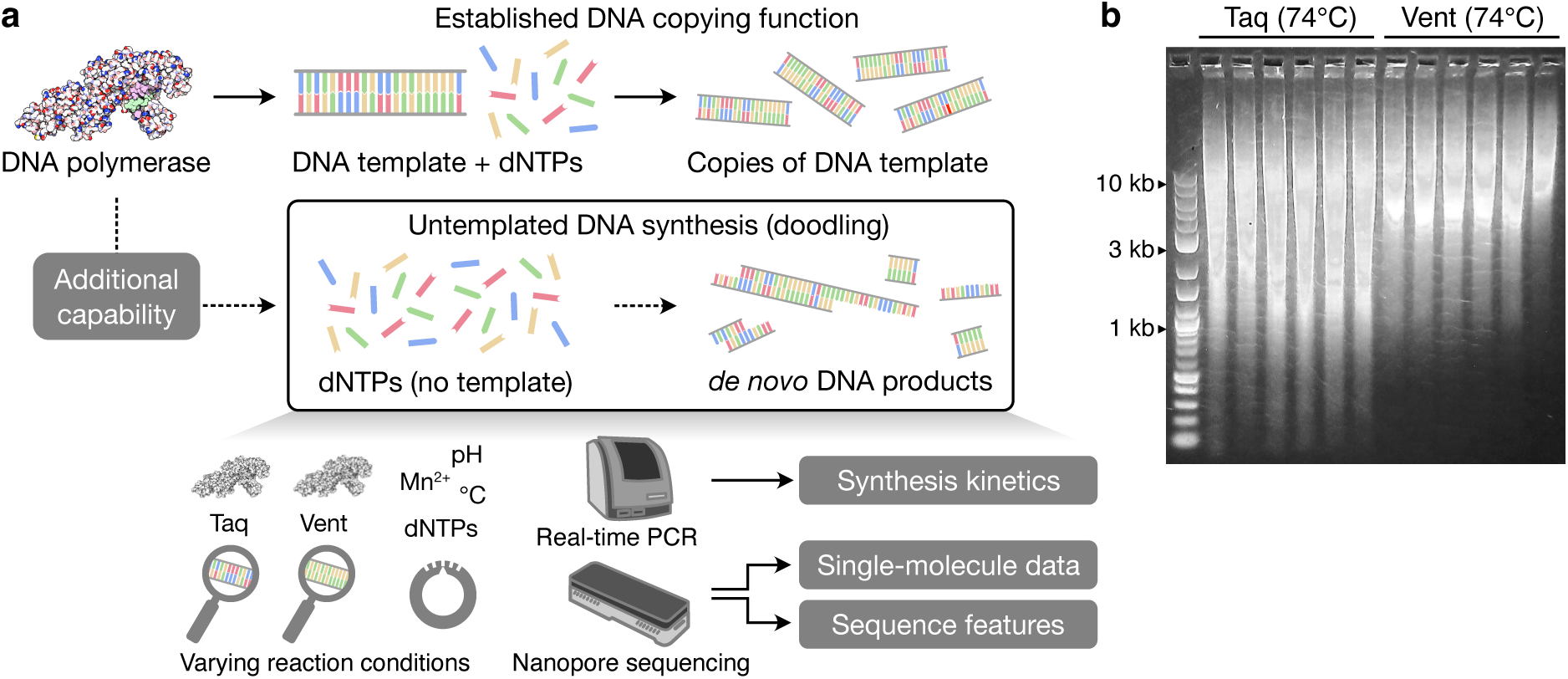
Methodology to study doodling by DNA polymerases. (**a**) Overview of the established copying function of DNA polymerases and the less well studies capability for them to perform untemplated DNA synthesis. In addition, methods used to study the process and the key data generated in this work are shown. (**b**) Varying length distribution of DNA doodled by Taq and Vent polymerases after 16 hour at 74°C with no DNA template present.

## Results

### Single molecule analysis of doodled DNA

We began by focusing on Taq and Vent polymerases that have been shown to be capable of untemplated DNA synthesis (**Figure 1b**) ^9,12^. Isothermal reactions were established using standard buffers for each polymerase at 65°C and 74°C, and run for 16 hours (**Methods**). We then performed nanopore sequencing of each reaction to characterise the sequences present in the pools of newly synthesized DNA fragments. To ensure the reads containing ‘doodled’ products were accurate, we trimmed sequencing adapters from the ends of each read, filtered out concatemer reads (namely those containing sequencing adapters within the read itself that likely occur due to ligation of separate fragments during the preparation of the sequencing libraries), and performed filtering based on the median Q score (≥13) and length (>65 nt) of each read to remove sequences that might correspond to DNA fragments introduced when preparing the sequencing libraries (i.e., adapters and barcodes) (**Methods**).

Analysis of the read length distributions showed substantive differences for each of the polymerases (**Figure 2**, top sections of each panel). Taq displayed a bimodal distribution with peaks at approximately 80 nt and 400 nt for reactions at 65°C, with a shift in the peak at 400 nt to 500 nt when the reaction temperature was increased to 74°C. This distribution also displayed a much longer tail (**Figure 2a,b**) with over 20% of reads >1000 nt compared to only 4–7% at 65°C (**Table 1**). For both temperatures, similar concentrations of synthesised DNA were measured (∼1000 ng/µL). In contrast, Vent had only a single peak in the read length distributions at ∼80 nt, with an exponential-type decay as read length increased that was similar across temperatures (**Figure 2c,d**). Additionally ∼10-fold less DNA was produced at the lower temperature and only small fractions of reads were >1000 nt long (≤6%) across both temperatures. This suggests Taq is more efficient at untemplated DNA synthesis and able to produce longer DNA fragments than Vent.

**Figure 2:**
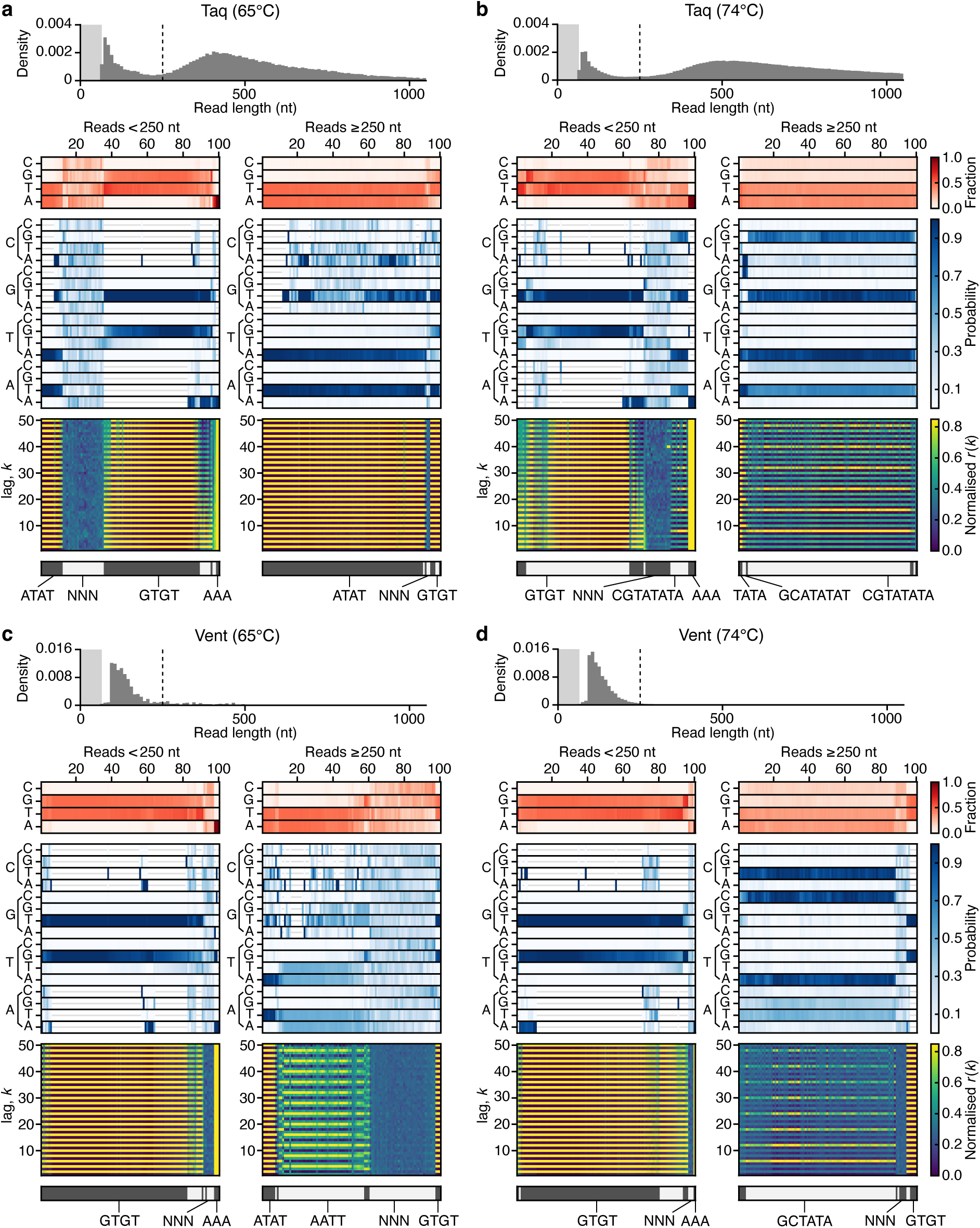
Single molecule analysis of doodling activity for Taq and Vent polymerases at different temperatures. (**a**) Taq at 65°C. (**b**) Taq at 74°C. (**c**) Vent at 65°C. (**d**) Vent at 74°C. In all panels, the top histogram shows the sequence length distribution with the lightly shaded region denoting the 0–65 nt range and dashed line denoting the 250 nt read length. Below this, heatmaps show for a random subset of reads smaller and larger than 250 nt (left and right plots, respectively) the following information (top–bottom): 1. Sequence composition (red heatmap), 2. Probability of transitioning from one base to another (blue heatmap), 3. Autocorrelation analysis capturing the similarity of the sequence compared to itself after varying nucleotide shifts/lag *k* (blue to yellow heatmap), and 4. the seven top clusters of reads (alternating light and dark grey) with key clusters having the sequence repeat they include below. Reads are displayed vertically and hierarchically clustered such that similar sequences are grouped.

**Table 1:**
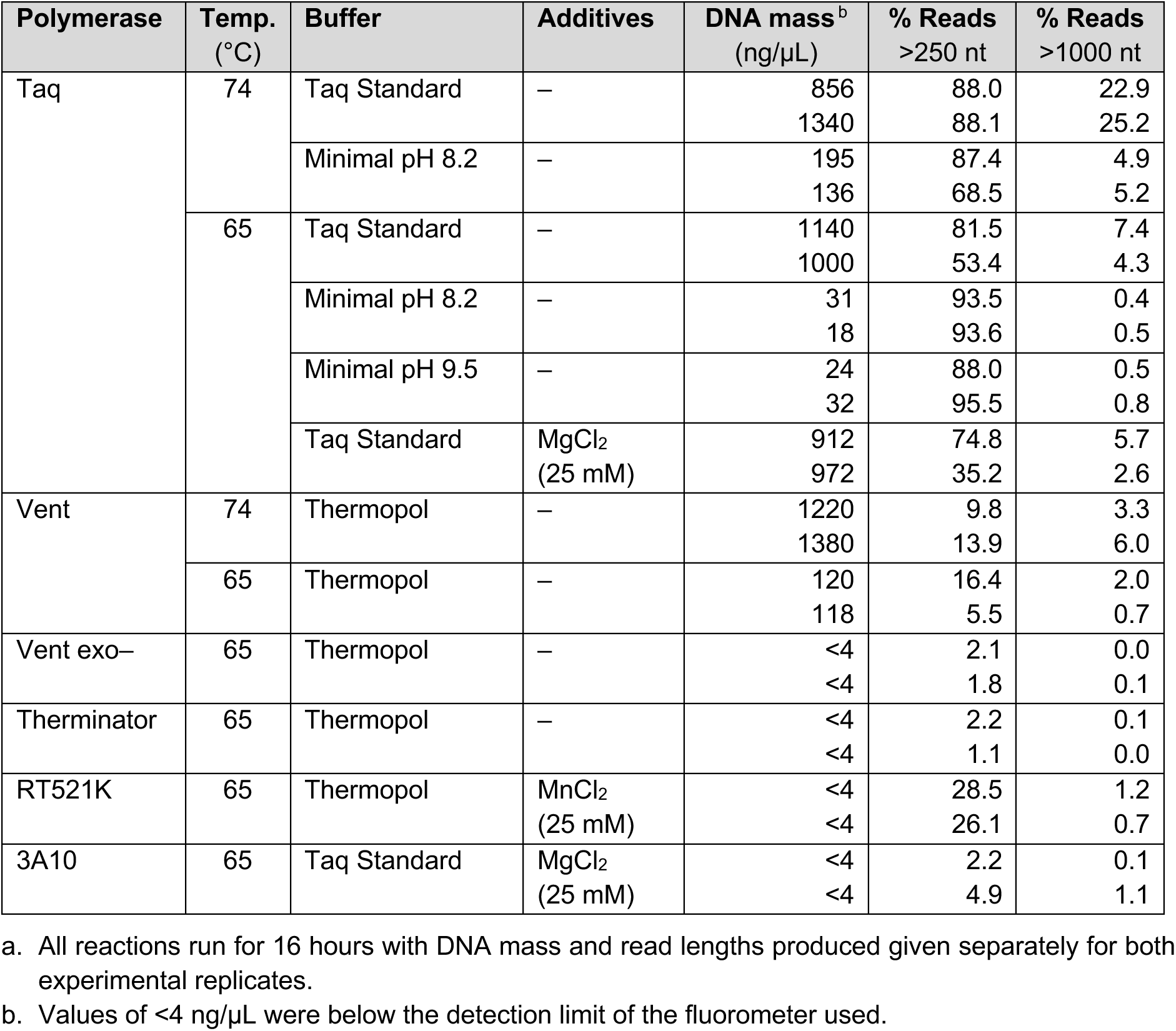
Doodling activities of DNA polymerases ^a^.

The nanopore sequencing additionally captured information regarding sequence composition and characteristic repetitive patterns (i.e., sequence motifs) within each DNA synthesized fragment. To characterize this dataset, we performed a number of linked analyses to generate visualisations of the DNA pools where each analysed read is denoted by a vertical column across all the plots (**Figure 2**, lower heatmaps). Analyses included assessing the base composition (red heatmaps) for each read, the transition probabilities from one base to another (blue heatmaps), and the sequence autocorrection to extract the scale (i.e., length/lag) of any repetitive sequences present (blue to yellow heatmaps). Each of these analyses was performed for a subset of reads shorter and longer than 250 nt and the results hierarchically clustered to ensure similar reads were grouped in the plots as vertical columns. The top seven clusters were highlighted to simplify their classification (alternating dark and light grey sections at the bottom of the panels). From these visualisations, it was possible to assess distinctive features of each doodled DNA sequence and to make comparisons between samples.

Using this approach, the Taq polymerase reactions at 65°C showed a majority of GT repeats (61%) for reads <250 nt, in addition to less frequent AT repeats (13%), poly-A sequences (4%), and random sequences (23%) (**Figure 2a**). In contrast, longer reads at this temperature were nearly exclusively AT repeats (91%), with only a small fraction of GT repeats and random sequences (9% in total). At 74°C, there was a distinctive shift in the sequence composition, with smaller reads retaining the dominance of GT repeats (71%), and longer reads demonstrating the emergence of CGTATATA repeats (93%) and a small fraction of GCATATAT (3%) repeats. The increased fraction of reads >1000 nt at 74°C, may relate to the fact that these longer repeats – when found in multiples of 3 or more – can efficiently self-extend by looping and base pairing of the complementary repeats and extension by Taq, using one or more of the single stranded repeats as a template ^5,14,15^.

For the Vent polymerase (**Figure 2c,d**), there was again a dominance of repetitive GT sequences (91–96%) and a small fraction of poly-A sequences (1–3%) for reads <250 nt across both temperatures. However, for longer reads >250 nt, there were large differences across temperatures as compared with Taq with Vent reactions at 65°C showing a small number of repetitive GT and AT sequences, instead being dominated by reads with AATT repeats (48%) and random sequences (37%) with near even coverage of bases (**Figure 2c**). This was substantially different to the AT repeat dominated Taq results (**Figure 2a**). Interestingly, the Vent reactions at 74°C produced a small fraction of longer reads containing GT repeats and random sequences (12% in total). However, the vast majority (88%) contained GCTATA repeats (**Figure 2d**). This again, differed from the longer CGTATATA repeat observed for the Taq reactions under similar conditions, and highlights potential biases in the affinities these polymerases have for the addition of specific nucleotides. At 74°C, the longer reads >250 nt demonstrate that Vent favours C → T, G → C, T → A and A → G or T, while Taq has a strong preference for C → G, G → T, T → A and A → T transitions. Furthermore, these preferences are influenced by temperature and appear to impact the efficiency with which DNA synthesis occurs (different transitions dominate the analysis of the short and long fragments).

These results were found to be robust for both polymerases tested and across different temperatures, with similar patterns observed in replicates (**Supplementary Figures 1 and 2**).

### Kinetics of doodling activity

Historically, the dynamics of doodling has principally been studied using time-sampling reactions and visualizing DNA products by gel electrophoresis (**Figure 3a**) ^17^. To more precisely quantify the doodling kinetics, we used real-time PCR to track the formation of DNA molecules synthesized under isothermal conditions. The experiments were run for 16 hours at 74°C for both the Taq and Vent polymerases. In both cases, we found that doodling occurred in two distinct stages (**Figure 3b**). Initially, a slow increase in synthesized DNA was observed. Then, at around 30 min, a transition to a second phase occurred with an approximately 3–3.5-fold increase in DNA synthesis rate. This rate was fixed (there was a linear increase in DNA mass) for a further ∼1.5 hours, after which the rate steadily decreased until DNA synthesis stopped ∼3 hours following the initiation of the reaction. This second stage appeared to capture a saturation in the ability for DNA synthesis, likely due to the depletion of free dNTPs, or production of DNA products that cannot be further extended. This behavior was consistent across replicates, with similar transition points and DNA synthesis rates (**Figure 3b**).

**Figure 3:**
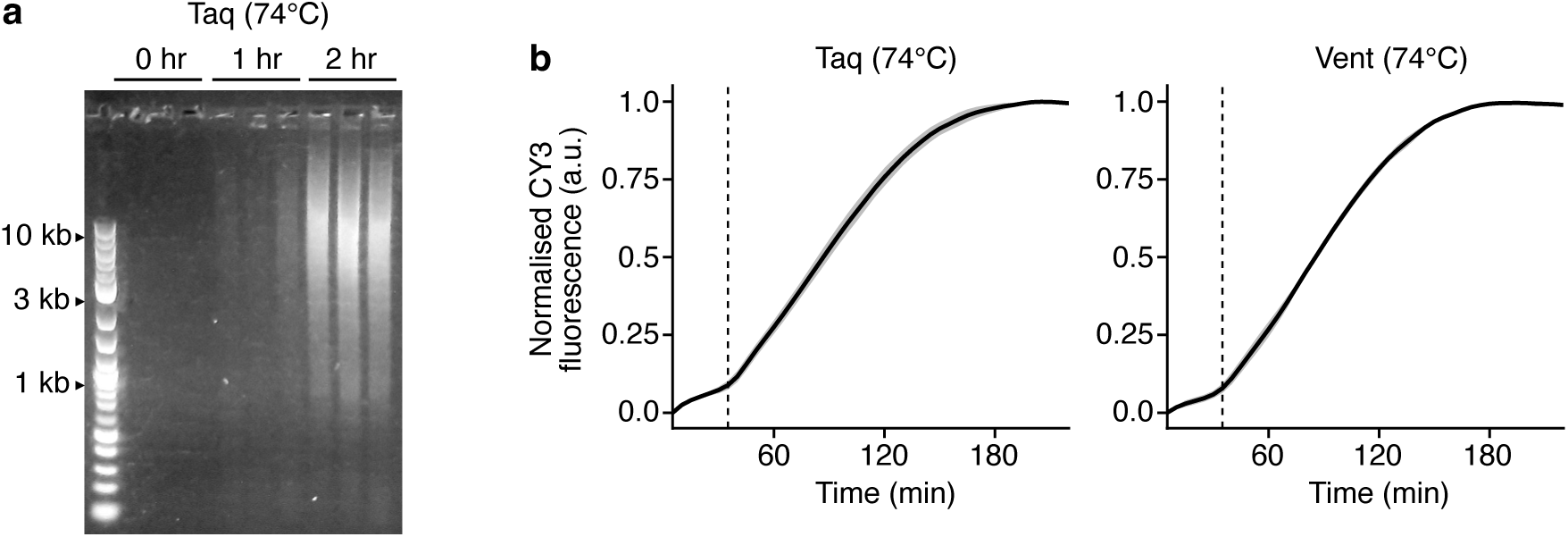
Kinetics of doodling activity for the Taq and Vent polymerases. (**a**) Emergence of doodled DNA products during an isothermal reaction at 74°C with Taq polymerase over 2 hours. (**b**) Normalized CY3 fluorescence for the Taq (left) and Vent (right) polymerases during the real-time PCR reaction. Data normalized by the maximum CY3 fluorescence of each sample. Solid black line shows the average of the 4 experimental replicates and grey shaded region denotes ±1 standard deviation. Thin dashed line denotes the 30 min point in the reaction.

This two-stage process is consistent with a previously suggested mechanism of untemplated DNA synthesis, whereby an initial stage allows for the generation of short random DNA sequences that may subsequently by chance discover repetitive sequences able to more efficiently self-replicate in a second stage ^5,14,15^ (also mentioned in the previous section). To our knowledge this is the first time that quantitative measurements of the duration and DNA synthesis rates for each stage have been made. Our results further support the role of self-replicating sequences in the generation of new genetic material.

### Influence of environmental factors

It has been shown that environmental factors such as temperature and buffer composition can influence the length and sequence composition of doodled DNA ^2,9,10,13^. While we have shown the important role that temperature plays in determining sequence composition (**Figure 2**), to further characterize this effect, we studied the doodling activity of Taq polymerase across a range of buffers. These included a minimal buffer with varying pH (8.2 and 9.5) and a standard Taq buffer including excessive amounts of MgCl_2_, which is known to stabilize the annealing of primers to incorrect template sites when used in PCR reactions and thereby decreases the specificity of the amplification process.

Similar to previous experiments, the results were shown to be highly reproducible for each condition, with similar patterns observed across experimental replicates (**Supplementary Figures 5–8**). A comparison of the sequence composition and repetitive motifs across different conditions showed that in most cases the same patterns were maintained. The only notable differences were for Taq at 65°C when using a minimal buffer at both pH 8.2 and 9.5, which showed a slight decrease (>2-fold) in the fraction of GC repeats for reads <250 nt (**Supplementary Figures 5 and 6**), and the more complex sequence repeats produced by Taq at 74°C in the minimal buffer at pH 8.2, which for one of the replicates (replicate 1) also displayed longer repetitive sequences at smaller (<250 nt) read lengths (**Supplementary Figure 8**).

The most notable differences observed were in the amount of DNA synthesized and the fraction of long (>1000 nt) DNA sequences produced (**Table 1**). Most of the reactions decreased the amount of DNA synthesized (apart from those at 65°C in the Taq Standard buffer with MgCl_2_), with the minimal buffer, leading to an ∼10-fold decrease at 74°C and ∼100-fold decrease at 65°C. Smaller drops were also seen in the percentage of long reads >1000 nt. Together, these results highlight the role that other environmental factors play in biasing the sequence composition and affecting the synthesis efficiency of the doodling process.

### Doodling activity of other DNAPs

Given the distinct sequence compositions of the doodled products derived using Taq and Vent polymerases, it appeared likely that other DNAPs may have unique biases in the untemplated sequences they generate. To further examine this, we considered two other DNAPs: Vent exo– to determine whether mutations in Vent that remove its proofreading exonuclease activity impact doodling, and Therminator as it was originally sourced from a distantly related species (*Thermococcus* species 9°N-7) and is known to incorporate modified substrates. In addition, we also studied two engineered DNAPs, RT521K and 3A10 ^18,19^. RT521K is an extensively engineered variant of Tgo polymerase ^18^ and 3A10 is a chimera of Tth and Taq polymerase generated by molecular breeding and compartmentalized self-replication (CSR) selection ^19^. We carried out 16 hour isothermal reactions and performed nanopore sequencing to characterize the sequences of the doodled DNA products (**Methods**).

Highly reproducible results were seen for all four polymerases (**Supplementary Figures 9–12**). For Vent exo–, we found large fractions of reads containing GT repeats (93% of reads <250 nt), a lack of the more complex AATT repeat sequences seen with standard Vent polymerase, and an increase in the faction of reads with near-random sequences (65% of reads >250 nt) (**Figure 4a**). Therminator displayed similar features, but contained a small number of longer reads (>250 nt) with more complex sequence repeats, including GCTATA, CGTATATA and AATT (3–8% total across replicates) (**Figure 4b**). For the engineered polymerases, we found that the short reads of RT521K were once again dominated by GT repeats, but virtually all longer reads >250 nt had random-like sequences (97%) (**Figure 4c**).

**Figure 4:**
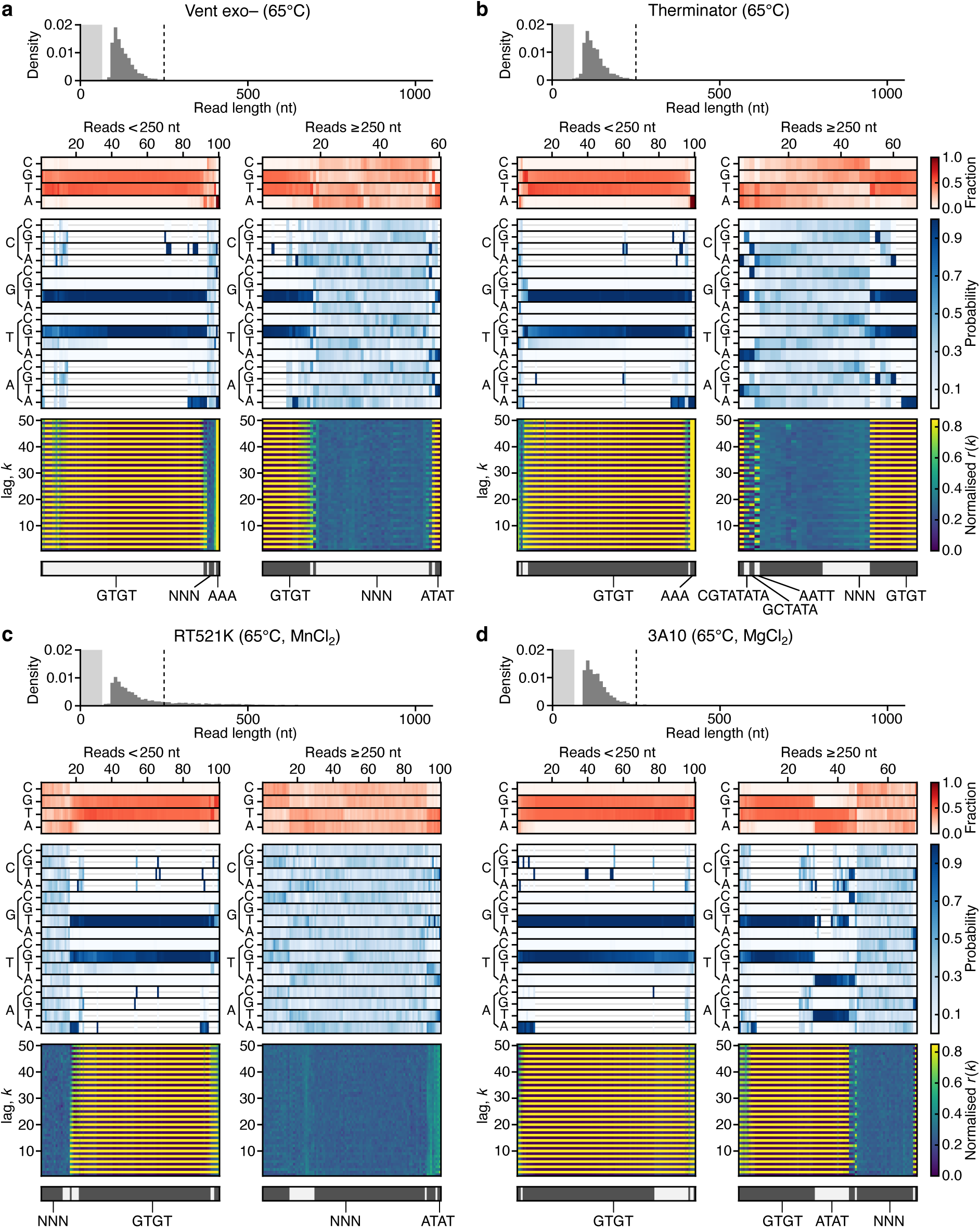
Single molecule analysis of doodling activity for diverse DNA polymerases. (**a**) Vent exo– at 65°C. (**b**) Therminator at 65°C. (**c**) RT521K at 65°C with 25mM MnCl_2_. (**d**) 3A10 at 65°C with 25mM MgCl_2_. In all panels, the top histogram shows the sequence length distribution with the lightly shaded region denoting the 0–65 nt range and dashed line denoting the 250 nt read length. Below this, heatmaps show for a random subset of reads smaller and larger than 250 nt (left and right plots, respectively) the following information (top–bottom): 1. Sequence composition (red heatmap), 2. Probability of transitioning from one base to another (blue heatmap), 3. Autocorrelation analysis capturing the similarity of the sequence compared to itself after varying nucleotide shifts/lag *k* (blue to yellow heatmap), and 4. the seven top clusters of reads (alternating light and dark grey) with key clusters having the sequence repeat they include below. Reads are displayed vertically and hierarchically clustered such that similar sequences are grouped.

This was evident from the near-equal fraction of bases and transition probabilities between bases. The 3A10 polymerase showed a similar dominance of random-like sequences, although to a lesser extent than RT521K (**Figure 4d**). When MnCl_2_ and MgCl_2_ were added to the buffers for the RT521K and 3A10 reactions, respectively, the probability of a shift to more random doodling became greater. For MnCl_2_, this is likely due to this additive promoting the misincorporation of nucleotides and destabilizing existing biases in base transitions ^20^.

The most distinctive difference of all the studied polymerases compared to Taq and Vent was the significantly reduced overall quantity of DNA synthesized per reaction (<4ng/µL in all cases) and greatly reduced fraction of reads >250 nt (**Table 1**).

### Analysis of repetitive sequence motifs

It is possible for a single DNA molecule to contain numerous different types of repetitive sequence throughout its length. These would not be captured by our previous autocorrelation analysis (**Figures 2 and 4**). To better characterize the full range of repetitive motifs present in the sequencing data, we performed an analysis in which we searched for a set of common motifs found across all the polymerases and conditions assayed. We then calculated the fraction of reads in which these repetitive motifs were found (**Figure 5**). It should be noted that this analysis is more sensitive than the autocorrelation analysis to single base differences, with exact motif sequences matches required and a minimum number of repeats (at least 7 consecutive repeats of 1–3 nt motifs, 6 consecutive repeats of 4 nt motifs, and 4 consecutive repeats of 6–8 nt motifs).

**Figure 5:**
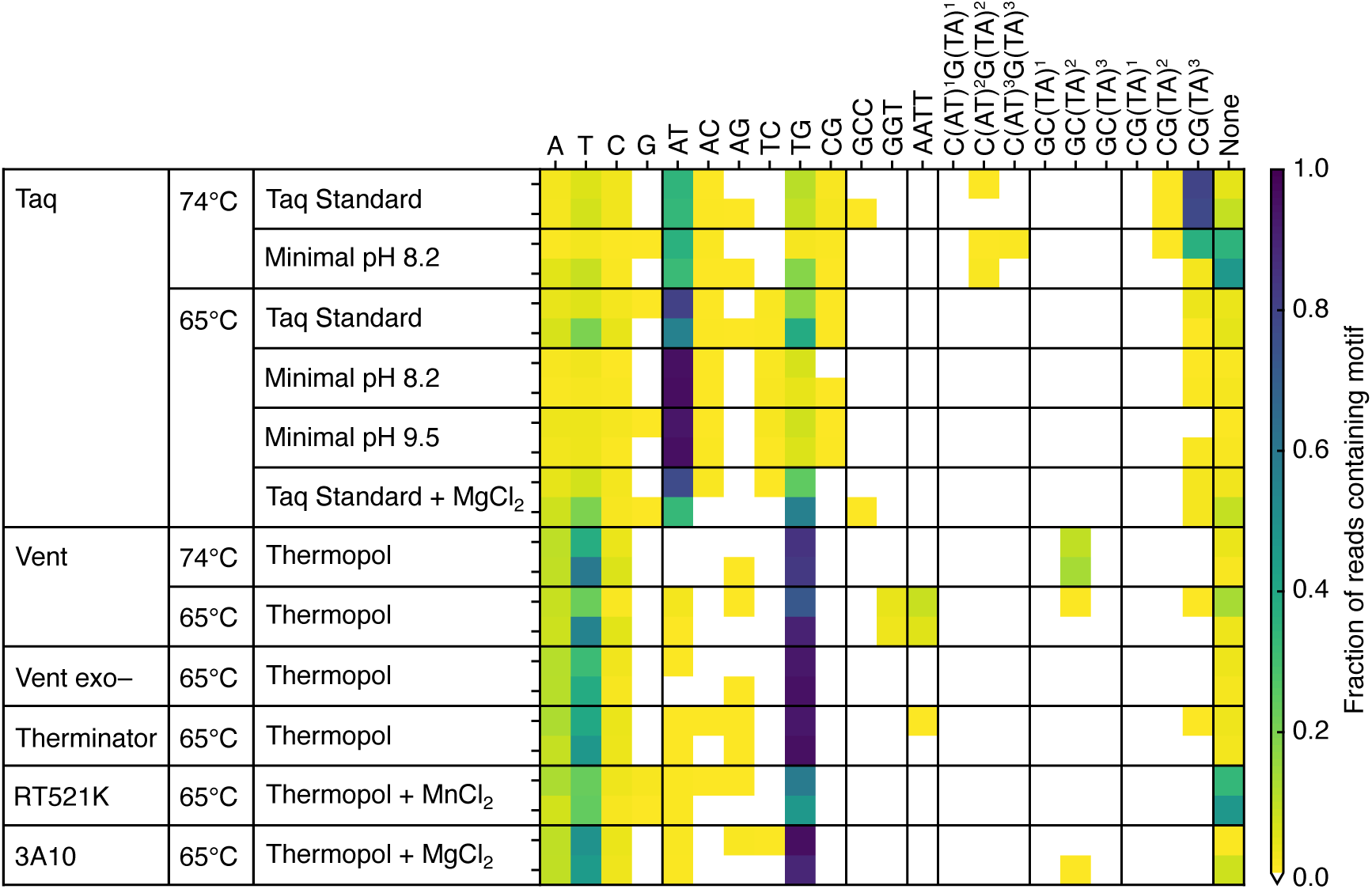
Composition of repetitive sequence motifs in doodled DNA. Heatmap shows the fraction of reads containing repetitive sequence motifs (columns) for different DNA polymerases and reaction conditions (rows). Results generated by randomly sampling 600 reads (>65 nt long) from the nanopore sequencing data and analyzing motif content of each separately. Data from the two experimental replicates is stacked with replicate 1 and 2 at the top and bottom, respectively. White areas of the heatmap denote motifs that were not present in any reads. To classify a read as containing a motif at least 7 consecutive repeats of 1–3 nt motifs, 6 consecutive repeats of 4 nt motifs, and 4 consecutive repeats of 6–8 nt motifs were required.

We observed compelling differences in the patterns of repetitive motif present for different polymerases, temperatures, and buffer compositions. One of the most striking features was the strong preference of AT repeats for Taq, and TG repeats for the other polymerases, as well as the synthesis of reads with few repetitive sequences for RT521K and Taq at 74°C with a minimal buffer at pH 8.2.

### Guiding sequence composition using dNTP availability

Having verified that experimental conditions like temperature and buffer composition could influence doodling behavior, we next sought to determine whether availability of dNTPs could be used to guide the sequence of doodled DNA, providing a potential means by which some degree of control could be obtained over the sequence composition of generated fragments.

In the first instance we carried out a set of additional real-time PCR experiments to determine whether the availability of only single types of dNTP, pairs of dNTP, or a set of three dNTP types (TCG) may influence the kinetics of doodling for the Taq and Vent polymerases (**Figure 6a**). We found that for single types of dNTP virtually no doodling was observed for both polymerases. In contrast, when two or more different types of dNTP were present, consistent normalized DNA synthesis curves were obtained that closely matched reactions in which all dNTPs were present for both Taq and Vent (**Figure 6a**). However, unlike reactions in which all dNTPs were present, most reactions with limited types of dNTP synthesized much less DNA <4 ng/µL compared to >100 ng/µL as compared with the situation when all dNTPs were present (**Supplementary Table 1**). This is unsurprising given the biases previously observed in the ability of DNAPs to incorporate specific dNTPs into a DNA molecule when a base of a particular type is present at the 3’ end.

**Figure 6:**
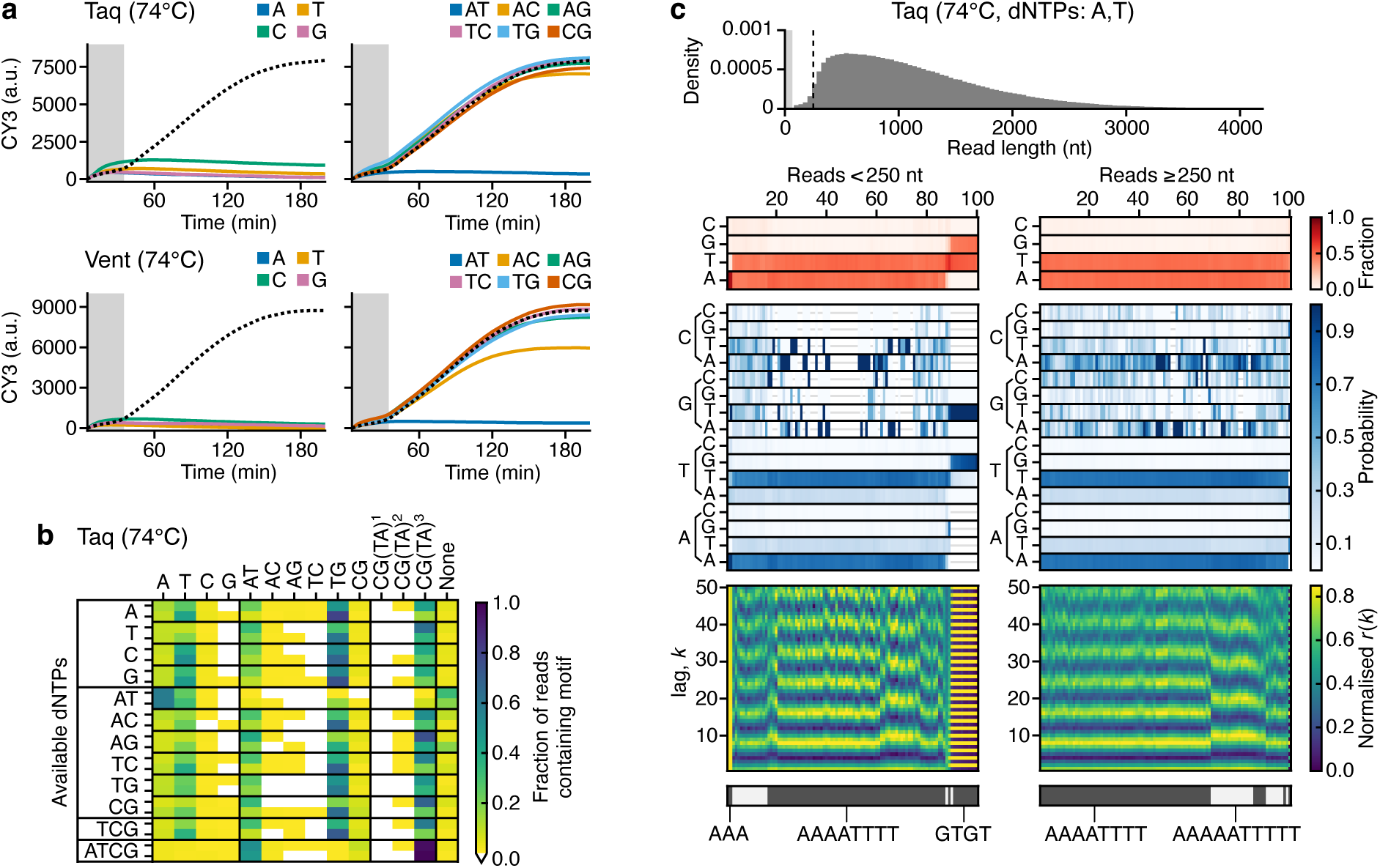
Guiding sequence composition of doodled DNA using dNTP availability. (**a**) CY3 fluorescence for the Taq (top) and Vent (bottom) polymerases during the real-time PCR reaction. (**b**) Heatmap shows the fraction of reads containing repetitive sequence motifs (columns) for Taq polymerase in Taq Standard buffer at 74°C for 16 hours, with different mixtures of dNTPs (rows). Results generated by randomly sampling 600 reads (>65 nt long) from the nanopore sequencing data and analyzing motif content of each separately. Data from the two experimental replicates is stacked with replicate 1 and 2 at the top and bottom, respectively. White areas of the heatmap denote motifs that were not present in any reads. To classify a read as containing a motif at least 7 consecutive repeats of 1–2 nt motifs, 6 consecutive repeats of 4 nt motifs, and 4 consecutive repeats of 6–8 nt motifs were required. (**c**) Sequence analysis for Taq at 74°C with only adenine and thymine present. Top histogram shows the sequence length distribution with the lightly shaded region denoting the 0–65 nt range and dashed line denoting the 250 nt read length. Below this, heatmaps show for a random subset of reads smaller and larger than 250 nt (left and right plots, respectively) the following information (top–bottom): 1. Sequence composition (red heatmap), 2. Probability of transitioning from one base to another (blue heatmap), 3. Autocorrelation analysis capturing the similarity of the sequence compared to itself after varying nucleotide shifts/lag *k* (blue to yellow heatmap), and 4. the seven top clusters of reads (alternating light and dark grey) with key clusters having the sequence repeat they include below. Reads are displayed vertically and hierarchically clustered such that similar sequences are grouped.

To determine whether the limited dNTP availability affects the composition of the sequences produced, we performed nanopore sequencing on each reaction. Analysis of the sequence motifs showed a bias for poly-T sequences and repeats of AT, TG and CGTATATA for all the single, double, and triple dNTP reactions, except for when only adenine and thymine were present (**Figure 6b**). It is unclear how DNA synthesis of bases not included in the reaction mix can take place, but could be due to contaminants in the individual dNTP mixes or polymerase preps used. A less likely cause could also be the nanopore sequencing library preparation, however, such sequences have not been seen previously and do not match any of the barcode or adapter sequences used. If this was contamination in the doodling reactions, this may help account for the small amounts of DNA synthesized in these reactions (**Supplementary Table 1**).

In cases where only adenine and thymine were present, we found, as expected, that sequences were nearly entirely composed of these bases (87% of reads <250 nt and 99% of reads >250 nt). Interestingly, short and long reads were dominated by AAAATTTT repeats in addition to AAAAATTTTT repeats at a lower frequency (**Figure 6c**). These patterns were robust and reproducible across experimental replicates (**Supplementary Figure 13**). Unlike the case with all other reactions with limited combinations of dNTPs where the majority of reads were <1000 nt (>80%), when only adenine and thymine were present >50% reads were long with a length >1000 nt. This suggests that these sequence motifs are more efficiently synthesized, likely due to their ability to self-replicate if the single-stranded DNA encoding them bends and base pairs with itself, allowing for standard extension and replication. This demonstrates the potential to modify the composition of doodled DNA, in some cases by tuning dNTP availability, and may offer a means for the synthesis of long DNA fragments where the specific sequence is less consequential, but where statistical composition must be maintained.

## Discussion

In this study we utilized long-read sequencing and real-time PCR to perform the most detailed analysis to date of untemplated DNA synthesis and obtained new insights into the sequence composition of the DNA fragment pools produced by natural and engineered DNAPs. Prior studies have characterized just a small number of separately cloned DNA sequences ^10^, making it impossible to understand the full diversity of sequences produced by untemplated synthesis activity and to identify rare sequences that may be of especial significance. Our results suggest that there may be distinct signatures of doodling activity resulting from biases in some of the base transitions and repetitive motifs. Further analysis of the occurrence of such signatures across genomes spanning all phylogeny could provide insights into a putative ongoing function of untemplated DNA synthesis, and genome evolution. Furthermore, building on previous work ^2,9,10,13^, we have shown that the sequence signatures observed are DNAP- dependent and may vary depending on environmental conditions and reaction buffer composition. The tuning of these parameters suggests a potential way in which the statistical features of doodled DNA sequences may be systematically manipulated and programmed. This has the potential to form the basis of a DNA synthesis method that could be disruptive to current oligonucleotide assembly methods.

Some efforts to exploit untemplated DNA synthesis by other enzymes has been made. Systems have been built that exploit the inherent sensitivity of terminal deoxynucleotidyl transferases (TdTs) to physiologically relevant signals like Co^2+^, Ca^2+^, Zn^2+^ and temperature to affect their template-independent extension as a means to encode environmental changes into DNA with a minute temporal resolution ^21^. Beyond direct synthesis, the evolution of biased DNA pools when a DNAP is present has also recently been investigated ^22^. Starting from a pool of purely adenine and thymine or guanine and cytosine DNA oligonucleotides (12 nt long) with a bias of purine or pyrimidine nucleotides, a *Bacillus stearothermophilus* strand displacing polymerase (Bst) was added and evolution of the pool monitored. It was found that initial biases in the pool disappeared as the oligonucleotides were extended and highly repetitive dimer and trimer motifs in rapidly extended sequences emerged. Analysis of these sequences revealed the importance of sequences supporting hairpin formation and reverse-complementary 3-mer periodic motifs. These findings provide some insights into the biophysical basis of the biases observed and may offer a way to better predict repetitive motifs that are likely to efficiently replicate if generated via a doodling reaction.

One major challenge we faced when working with pools of doodled DNA is the extreme length of some fragments. While the read length distributions we report give some idea of differences in fragment lengths between experiments, it is likely that extremely long DNA fragments are not captured during sequence analysis due to fragmentation by shearing during preparation of the sequencing libraries. We have used barcoding to enable the multiplexing of samples, but this introduces additional liquid handling steps that may have increased the extent and frequency of DNA shearing events. We have carried out some preliminary sequencing experiments where the barcoding stage was omitted, and found that there was a shift in the read length distributions towards longer reads. In some cases reads >23 kb were recorded (maximum lengths of 23,508 bp and 85,364 bp for Taq and Vent, respectively) (**Supplementary Figure 14**). The distributions presented here are therefore likely to represent the lower boundary of the actual DNA fragment lengths generated by doodling. Developing new library preparation methods that mitigate shearing comprises one important future direction of this work ^23,24^.

The ability to use doodling for the production of extremely long DNA fragments (potentially >100 kb) differentiates it from existing technologies that are typically based on relatively slow cyclic processes to enable the syntheses of specific DNA sequences ^25^. However, our results show that while the native doodling capabilities of DNAPs vary in response to environmental changes, these changes are limited and cannot yet be precisely controlled to support the synthesis of specific DNA sequences. Developments in long-read sequencing through the engineering of polymerases and protein nanopores and their interfacing with electrical and photonic systems have demonstrated the leaps that can be made to rapidly harness and repurpose biological functions ^26–31^. Therefore, an interesting future direction would be to explore whether the interfacing of DNAPs with electronic systems might facilitate the rapid modulation of environmental or catalytic biases in the DNAP and greater control over the doodling process and composition of the DNA pools synthesised.

If sufficient improvements to the control of the doodling process can be made, it may be possible to rapidly and efficiently produce long ssDNA fragments of defined composition. This would have significant implications for a broad range of biotechnological applications. In the near term, the ability to accurately synthesize long DNA molecules rapidly, and at low cost, will have substantive utility in accelerating large-scale genetic circuit construction ^32^ and could act as a novel substrate for material sciences, e.g., DNA-based “glues” ^25,33,34^. Furthermore, the ability for the sequences generated to dynamically vary in response to specific environmental conditions provides a means for recording events as sequences in stable DNA molecules (e.g., similar to Bhan *et al.* ^21^). Over the longer term, this work builds on ongoing efforts to establish a foundation for exploring novel, biologically inspired approaches to DNA synthesis. Compared to existing approaches (such as DNA synthesis using phosphoramidite chemistry) these would have a less significant environmental imprint, and would be faster, more efficient, and have the potential to generate DNA sequences of virtually any length ^25,35^. This type of technology is exactly what will be required for biological engineering as it transitions from single-gene interventions to system-level genome design and construction, and is broadly utilized as a predictive engineering material ^36,37^. This work supports these efforts and further demonstrates the value of long read sequencing when attempting to unpick complex genetic processes ^30,38,39^.

## Methods

### In vitro reactions

All doodling reactions were performed in thin wall PCR tubes or 96-well PCR plates, using reaction volumes of 100 µL with 200 µM of dNTPs (New England Biolabs, N0447S). Reactions were prepared on ice in a PCR hood before being transferred to a preheated thermocycler (Bio-Rad, C1000). Nuclease-free water (Omega Biotek, PD092) was used to prepare all reactions. Reactions were run for 16 hours at 64°C or 75°C, unless stated otherwise. Taq reactions were performed in a 1X Taq Standard Buffer (New England Biolabs, B9014) containing 10 mM Tris-HCl, 50 mM KCl, 1.5 mM MgCl_2_, and 5 units of polymerase (New England Biolabs M0273). Vent reactions were performed in 1X Thermopol buffer (New England Biolabs, B9004) containing 20 mM Tris-HCl, 10 mM (NH_4_)_2_SO_4_, 10 mM KCl, 2 mM MgSO_4_, 0.1% Triton X-100, and 2 units of polymerase (New England Biolabs M0254). Vent exo– (New England Biolabs, M0257) and Therminator (New England Biolabs, M0261) polymerase reactions were set up similarly to those for Vent. RT521K polymerase reactions were performed in 1X Thermopol buffer supplemented with 25 Mm MnCl_2_. 3A10 polymerase reactions were performed in 1X Taq standard buffer supplemented with 25 mM MgCl_2_. For RT521K and 3A19 polymerase reaction, 1 µL (5 U/µL) of polymerase was used. 1X minimal buffer was formulated as 10 mM Tris-HCl, 50 mM KCl, 2.5 mM MgCl_2_ and adjusted to pH 8.2 at 25°C or pH 9.5 at 25°C.

### Gel electrophoresis

Gel electrophoresis was performed for DNA length quantification of reaction products. Gels were prepared using 1% agarose in 30–50 mL of 1X TAE running buffer (40 mM Tris-acetate, 1 mM EDTA). Gels were stained by adding 1X nucleic acid stain Gel Green (Biotium, 41005) to the gel solution prior to casting. 50 µL of sample was loaded into each lane of the gel, using 10 µL purple loading dye (New England Biolabs, B7025S). The DNA 1 kb+ ladder (New England Biolabs, N3200) was used as a reference. Gels were run at 80V for 30–60 minutes (Bio-Rad, PowerPac Basic).

### Nanopore DNA sequencing

DNA was prepared for nanopore sequencing following the ligation sequencing of amplicons protocol using Oxford Nanopore Technologies (ONT) kit SQK-LSK110 directly, or the native barcoding protocol using ONT kits SQK-LSK110 and EXP-NBD104. DNA quantification was performed using a Qubit dsDNA broad range DNA quantification kit and a Qubit 4 fluorometer (Invitrogen, Q33238). For barcoding, equal volumes of each barcoded sample were pooled and 6–60 ng of DNA was loaded into each flow cell (ONT, FLO-MIN106). Sequencing was performed using a MinION Mk1B device and the MinKNOW software version 21.11.7.

### Real-time PCR

Real-time PCR was used to monitor the synthesis of doodled DNA products during a reaction. For the Taq polymerase, 100 µL reactions were prepared on ice in 96-well plates containing 84 µL nuclease free water (Omega Biotek, PD092), 10 µL of 10X Taq standard buffer (New England Biolabs, B9014), 2 µL of 10mM dNTPs (complete mixtures of all dNTPs, New England Biolabs, N0447S; or combinations of single types of dNTP supplied as a separate set, New England Biolabs, N0446S), 1 µL Taq polymerase (2 units final concentration), 2 µL ROX solution (ThermoFisher Scientific, A46109), and 1 µL SYBR Green dye (ThermoFisher Scientific, A46109). For the Vent polymerase, identical reactions were prepared, however, the 10X Taq standard buffer was replaced with a 10X Thermopol buffer (New England Biolabs, B9004). Assembled reactions were then run in a preheated qPCR machine (Agilent Technologies, Stratagene MXP3005P) with a fixed temperature of 74°C and a cycle time of 5 min for a total of 16 hours. Measurements of fluorescence (CY3 and ROX) were taken each cycle.

### Data analysis

Raw sequencing data in FAST5 format was basecalled using guppy version 6.3.8 and the configuration file ‘dna_r9.4.1_450bps_hac.cfg’. The resultant FASTQ files were then cleaned by trimming any adapter or barcode sequences found at the start or end of each read and reads with barcodes found internally were removed. Unless otherwise stated, we also filtered reads based on length >65 nt and Q-score ≥13 to enable more accurate inference of statistical and repetitive features of the sequences. Autocorrelation analyses were performed by calculating a discrete and normalized string-based autocorrelation function *r*(*k*), where *k* is the lag (shift in sequence). We assume that only perfectly matching elements (bases) are included in the calculation with the final value normalized by sequence length to account for the finite nature of the input sequence. This was implemented as a custom function in Julia (**Supplementary Data 1**). All analysis scripts were developed and run using Julia version 1.10.2 and the following packages: BioAlignments version 3.1.0, BioSequences version 3.1.6, Clustering version 0.15.7, CSV version 0.10.14, DataFrames version 1.6.1, Dates version 1.10.0, Distances version 0.10.11, FASTX version 2.1.4, Glob version 1.3.1, LinearAlgebra version 1.10.0, Random version 1.10.0, and Statistics version 1.10.0. Plots were generated using Julia and the CairoMakie version 0.12.3 package and final figures composited in Affinity Designer version 2.5.3.

## Data Availability

Sequencing data, real-time PCR data and analysis scripts are available from Zenodo at: https://doi.org/10.5281/zenodo.13388199

## Acknowledgements

This research was supported by Replay Holdings Inc., a Royal Society University Research Fellowship grant URF\R\221008 (T.E.G.), a Turing Fellowship from The Alan Turing Institute under the Engineering and Physical Sciences Research Council (EPSRC) grant EP/N510129/1 (T.E.G.), BrisEngBio, a UKRI Engineering Biology Transition Award grant BB/W013959/1 (T.E.G.), and by the Medical Research Council (MRC) as part of United Kingdom Research and Innovation (also known as UK Research and Innovation (UKRI)) MRC program grant MC_U105178804 (P.H.) We would also like to thank the Replay Genome Writing Team for insightful discussions and feedback on the research. For the purpose of open access, the MRC Laboratory of Molecular Biology has applied a CC BY public copyright license to any Author Accepted Manuscript version arising.

## Author contributions

A.W., A.H., T.E.G. and G.L. conceived the project. T.E.G., S.D.C., A.W. and G.L. designed the experiments. P.H. provided the 3A10 and RT521K engineered DNA polymerases for testing. S.D.C. performed all experiments. T.E.G. and S.D.C. carried out all the analyses with input from G.L., B.T.R. and I.D.W.S. T.E.G. supervised all experiments and wrote the initial manuscript. All authors contributed to the interpretation of the results and editing of the manuscript.

