## Supplementary Information for "Analysis and control of untemplated DNA polymerase activity for guided synthesis of kilobase-scale DNA sequences"


| **Supplementary Figures** | | **Page** |
|---|---|---|
| Supplementary Figure 1: | Sequence analysis of Taq (65°C, Taq Standard Buffer) | 2 |
| Supplementary Figure 2: | Sequence analysis of Taq (74°C, Taq Standard Buffer) | 3 |
| Supplementary Figure 3: | Sequence analysis of Vent (65°C, Thermopol Buffer) | 4 |
| Supplementary Figure 4: | Sequence analysis of Vent (74°C, Thermopol Buffer) | 5 |
| Supplementary Figure 5: | Sequence analysis of Taq (65°C, Minimal pH 8.2 Buffer) | 6 |
| Supplementary Figure 6: | Sequence analysis of Taq (65°C, Minimal pH 9.5 Buffer) | 7 |
| Supplementary Figure 7: | Sequence analysis of Taq (65°C, Taq Standard Buffer, 25 mM $MgCl_2$) | 8 |
| Supplementary Figure 8: | Sequence analysis of Taq (74°C, Minimal pH 8.2 Buffer) | 9 |
| Supplementary Figure 9: | Sequence analysis of Vent exo– (65°C, Thermopol Buffer) | 10 |
| Supplementary Figure 10: | Sequence analysis of Therminator (65°C, Thermopol Buffer) | 11 |
| Supplementary Figure 11: | Sequence analysis of RT521K (65°C, Thermopol Buffer, 25 mM $MnCl_2$) | 12 |
| Supplementary Figure 12: | Sequence analysis of 3A10 (65°C, Taq Standard Buffer, 25 mM $MgCl_2$) | 13 |
| Supplementary Figure 13: | Sequence analysis of Taq (65°C, Taq Standard Buffer, only adenine and thymine provided) | 14 |
| Supplementary Figure 14: | Read length distributions from non-barcoded sequencing runs | 15 |
| **Supplementary Tables** | | |
| Supplementary Table 1: | Doodling activity of Taq with differing dNTP availability | 16 |

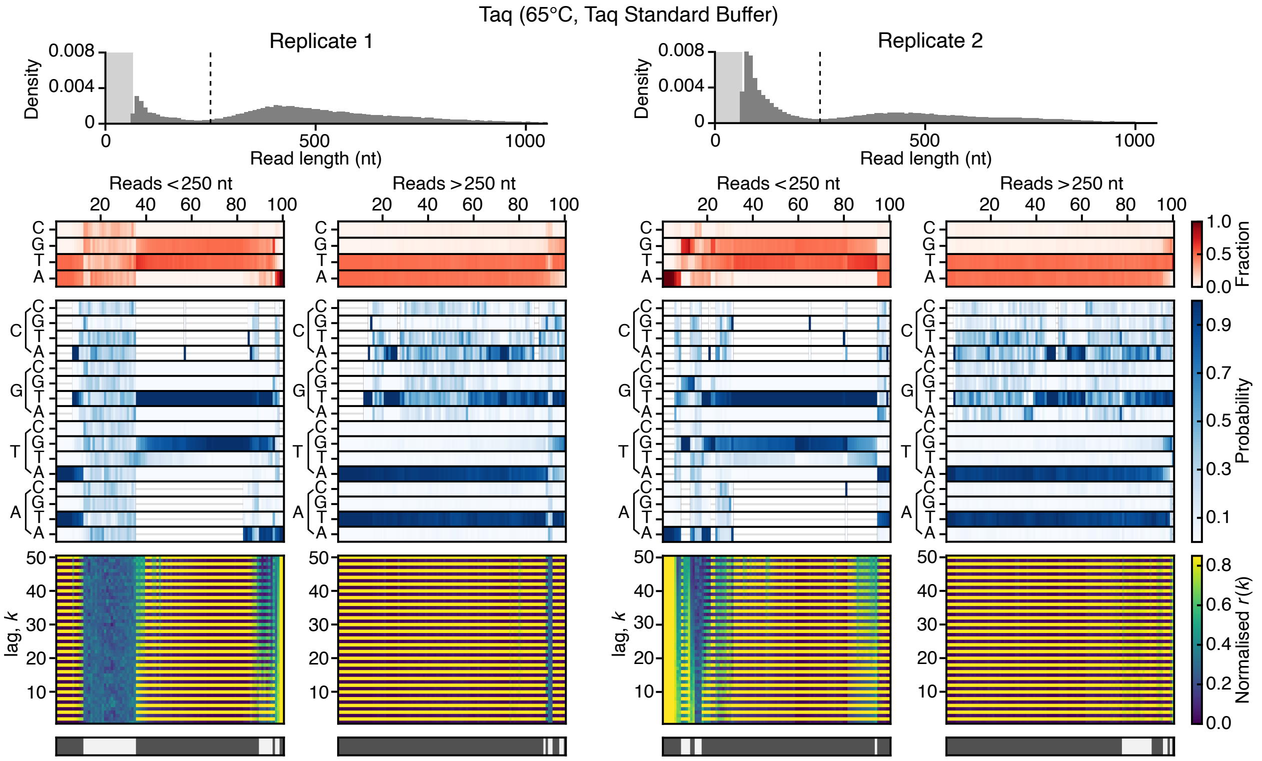


**Supplementary Figure 1: Sequence analysis of Taq (65°C, Taq Standard Buffer).** The top histograms show the sequence length distribution with the lightly shaded region denoting the 0–65 nt range and dashed line denoting the 250 nt read length. Below this, heatmaps show for a random subset of reads smaller and larger than 250 nt (left and right plots, respectively) the following information (top–bottom): 1. Sequence composition (red heatmap), 2. Probability of transitioning from one base to another (blue heatmap), 3. Autocorrelation analysis capturing the similarity of the sequence compared to itself after varying nucleotide shifts/lag *k* (blue to yellow heatmap), and 4. the seven top clusters of reads (alternating dark and light grey). Reads are displayed vertically and hierarchically clustered such that similar sequences are grouped.

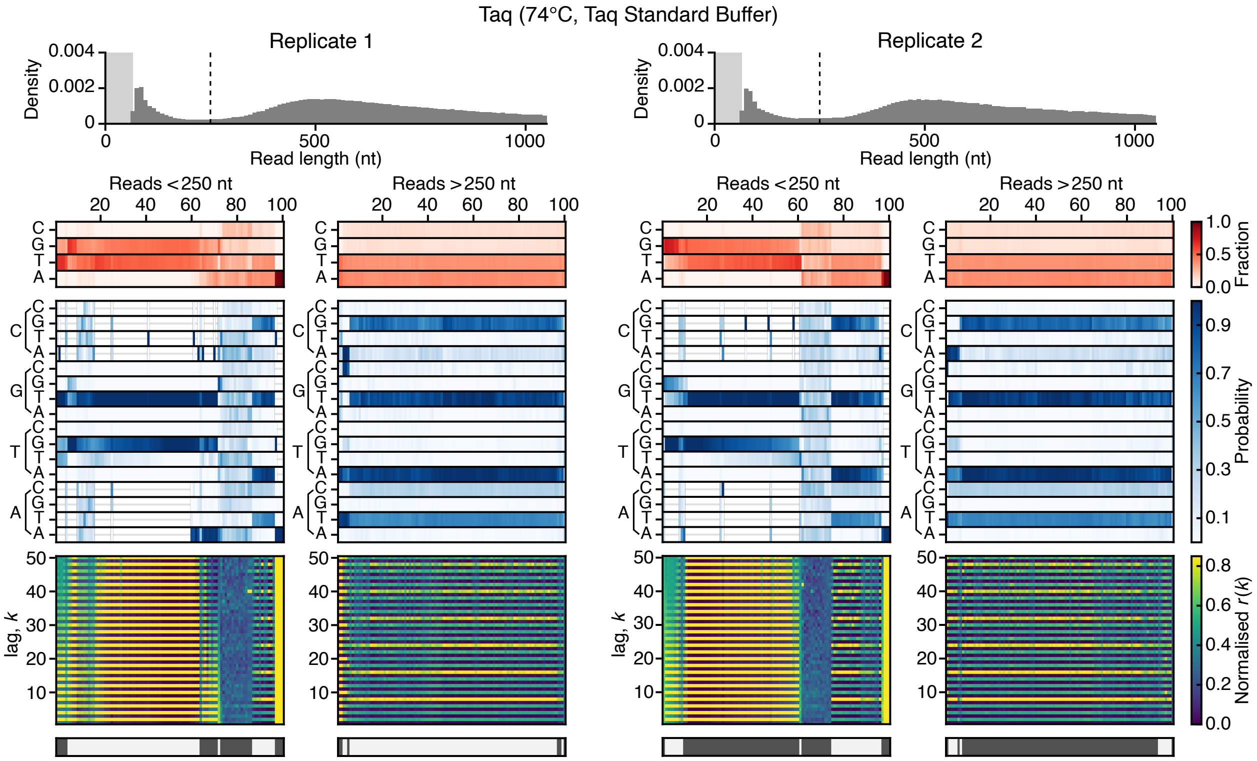


**Supplementary Figure 2: Sequence analysis of Taq (74°C, Taq Standard Buffer).** The top histograms show the sequence length distribution with the lightly shaded region denoting the 0–65 nt range and dashed line denoting the 250 nt read length. Below this, heatmaps show for a random subset of reads smaller and larger than 250 nt (left and right plots, respectively) the following information (top–bottom): 1. Sequence composition (red heatmap), 2. Probability of transitioning from one base to another (blue heatmap), 3. Autocorrelation analysis capturing the similarity of the sequence compared to itself after varying nucleotide shifts/lag *k* (blue to yellow heatmap), and 4. the seven top clusters of reads (alternating dark and light grey). Reads are displayed vertically and hierarchically clustered such that similar sequences are grouped.

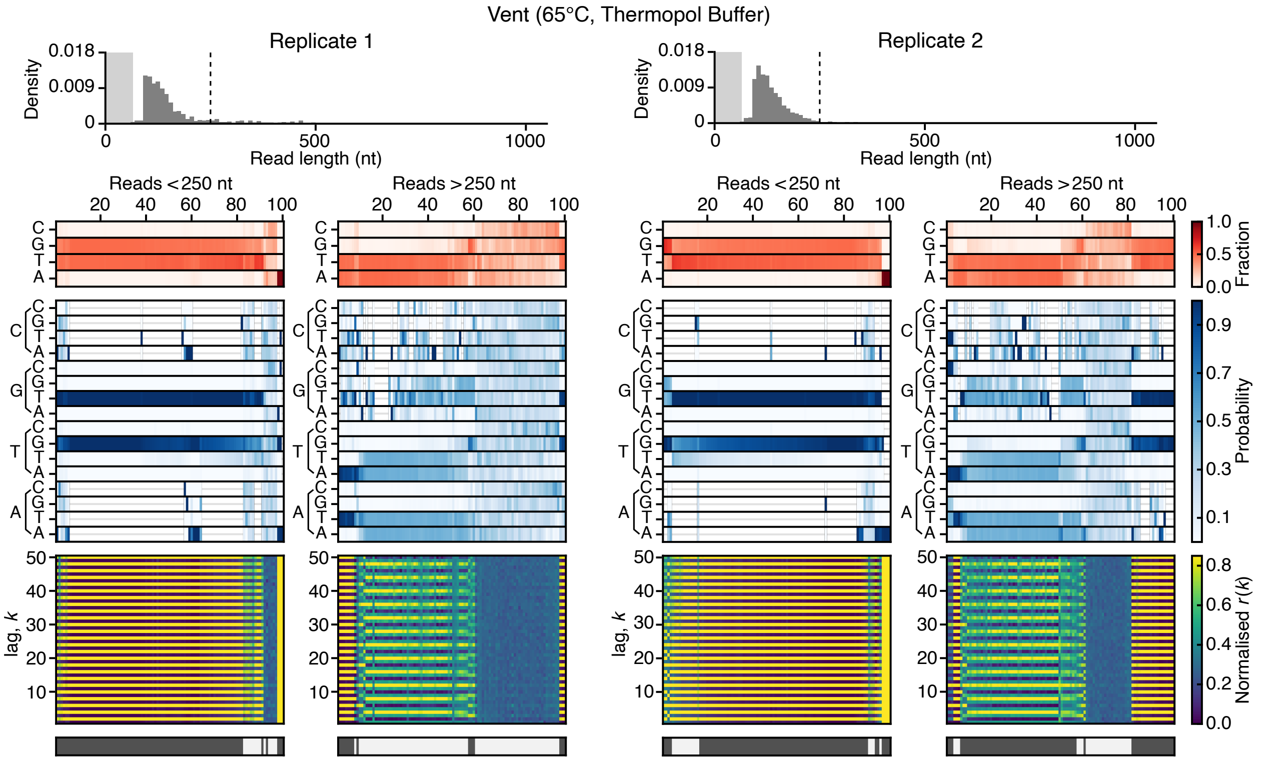


**Supplementary Figure 3: Sequence analysis of Vent (65°C, Thermopol Buffer).** The top histograms show the sequence length distribution with the lightly shaded region denoting the 0–65 nt range and dashed line denoting the 250 nt read length. Below this, heatmaps show for a random subset of reads smaller and larger than 250 nt (left and right plots, respectively) the following information (top–bottom): 1. Sequence composition (red heatmap), 2. Probability of transitioning from one base to another (blue heatmap), 3. Autocorrelation analysis capturing the similarity of the sequence compared to itself after varying nucleotide shifts/lag $k$ (blue to yellow heatmap), and 4. the seven top clusters of reads (alternating dark and light grey). Reads are displayed vertically and hierarchically clustered such that similar sequences are grouped.

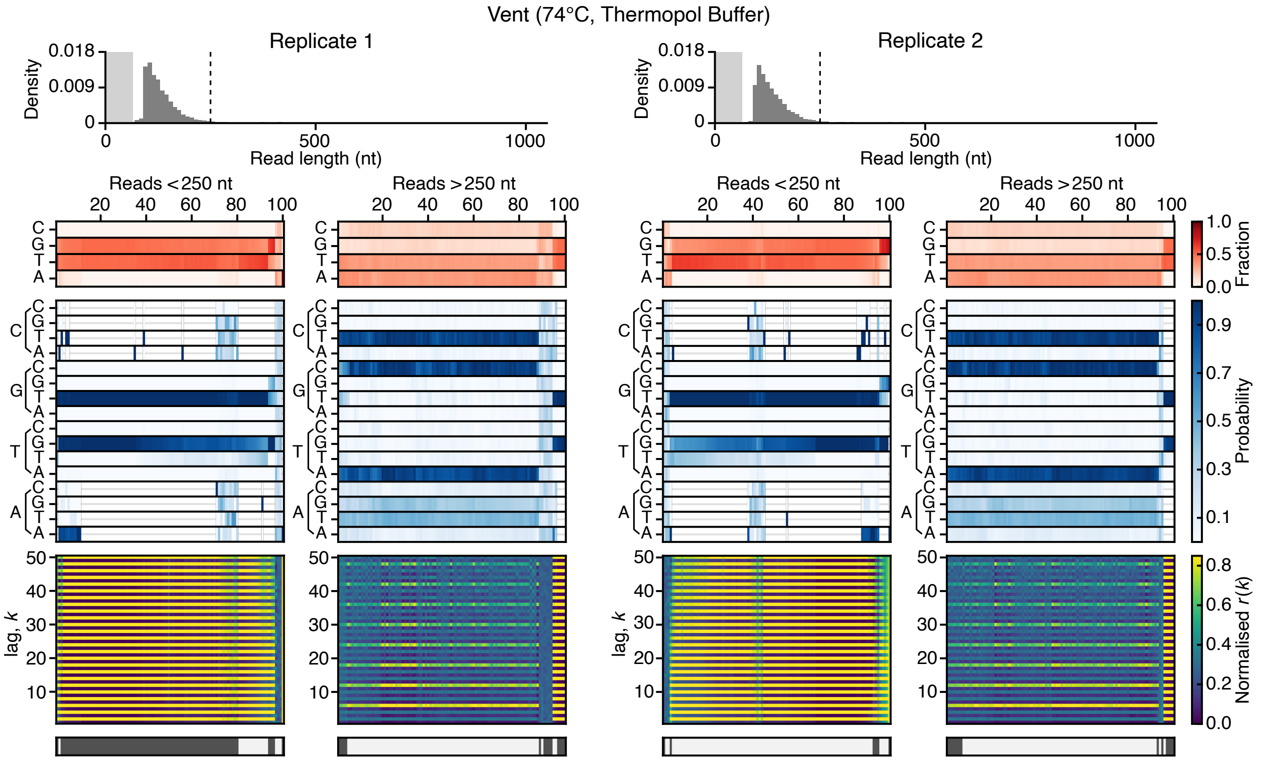


**Supplementary Figure 4: Sequence analysis of Vent (74°C, Thermopol Buffer).** The top histograms show the sequence length distribution with the lightly shaded region denoting the 0–65 nt range and dashed line denoting the 250 nt read length. Below this, heatmaps show for a random subset of reads smaller and larger than 250 nt (left and right plots, respectively) the following information (top–bottom): 1. Sequence composition (red heatmap), 2. Probability of transitioning from one base to another (blue heatmap), 3. Autocorrelation analysis capturing the similarity of the sequence compared to itself after varying nucleotide shifts/lag *k* (blue to yellow heatmap), and 4. the seven top clusters of reads (alternating dark and light grey). Reads are displayed vertically and hierarchically clustered such that similar sequences are grouped.

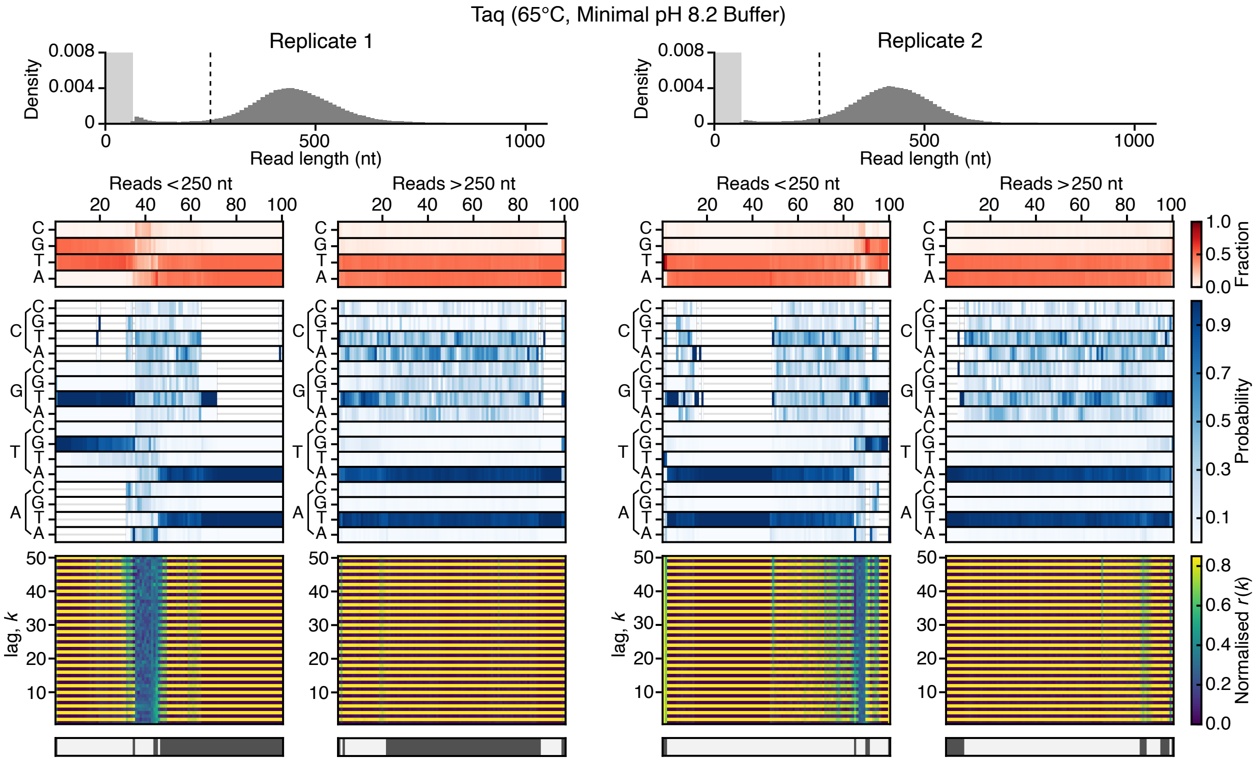


**Supplementary Figure 5: Sequence analysis of Taq (65°C, Minimal pH 8.2 Buffer).** The top histograms show the sequence length distribution with the lightly shaded region denoting the 0–65 nt range and dashed line denoting the 250 nt read length. Below this, heatmaps show for a random subset of reads smaller and larger than 250 nt (left and right plots, respectively) the following information (top–bottom): 1. Sequence composition (red heatmap), 2. Probability of transitioning from one base to another (blue heatmap), 3. Autocorrelation analysis capturing the similarity of the sequence compared to itself after varying nucleotide shifts/lag *k* (blue to yellow heatmap), and 4. the seven top clusters of reads (alternating dark and light grey). Reads are displayed vertically and hierarchically clustered such that similar sequences are grouped.

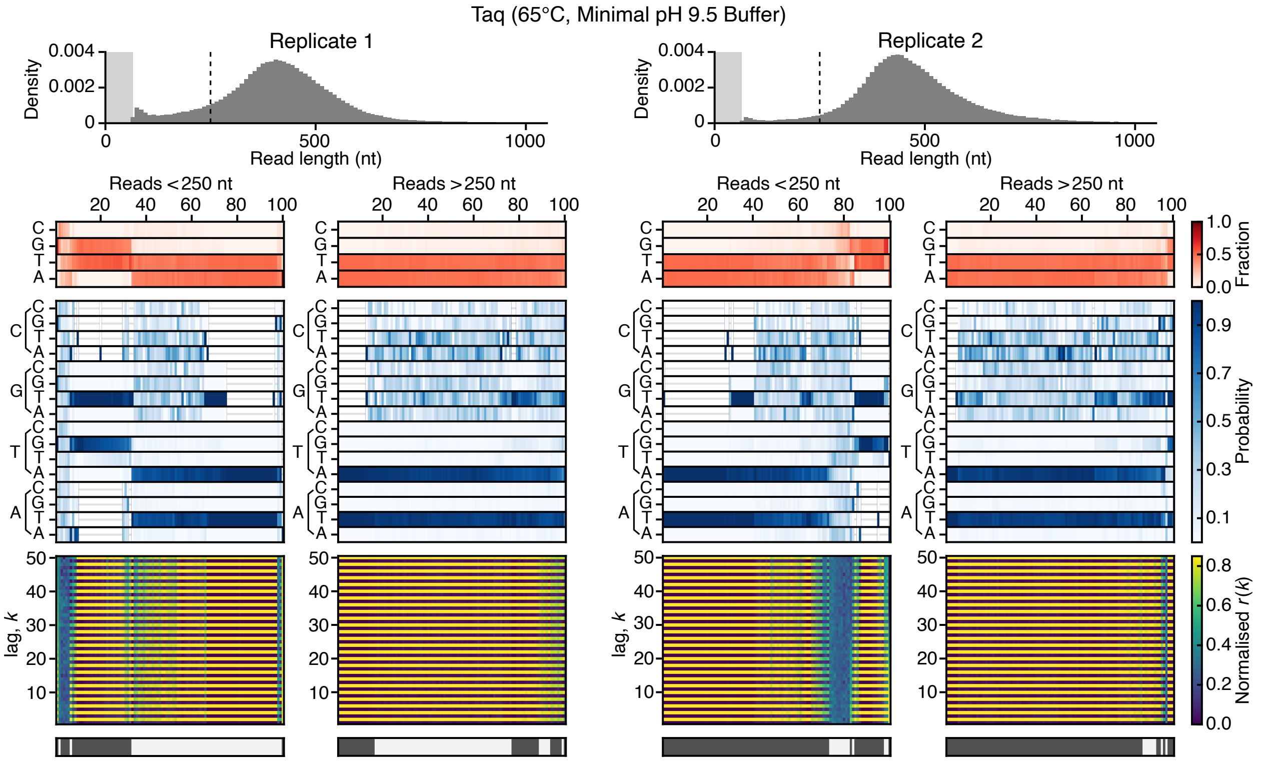


**Supplementary Figure 6: Sequence analysis of Taq (65°C, Minimal pH 9.5 Buffer).** The top histograms show the sequence length distribution with the lightly shaded region denoting the 0–65 nt range and dashed line denoting the 250 nt read length. Below this, heatmaps show for a random subset of reads smaller and larger than 250 nt (left and right plots, respectively) the following information (top–bottom): 1. Sequence composition (red heatmap), 2. Probability of transitioning from one base to another (blue heatmap), 3. Autocorrelation analysis capturing the similarity of the sequence compared to itself after varying nucleotide shifts/lag $k$ (blue to yellow heatmap), and 4. the seven top clusters of reads (alternating dark and light grey). Reads are displayed vertically and hierarchically clustered such that similar sequences are grouped.

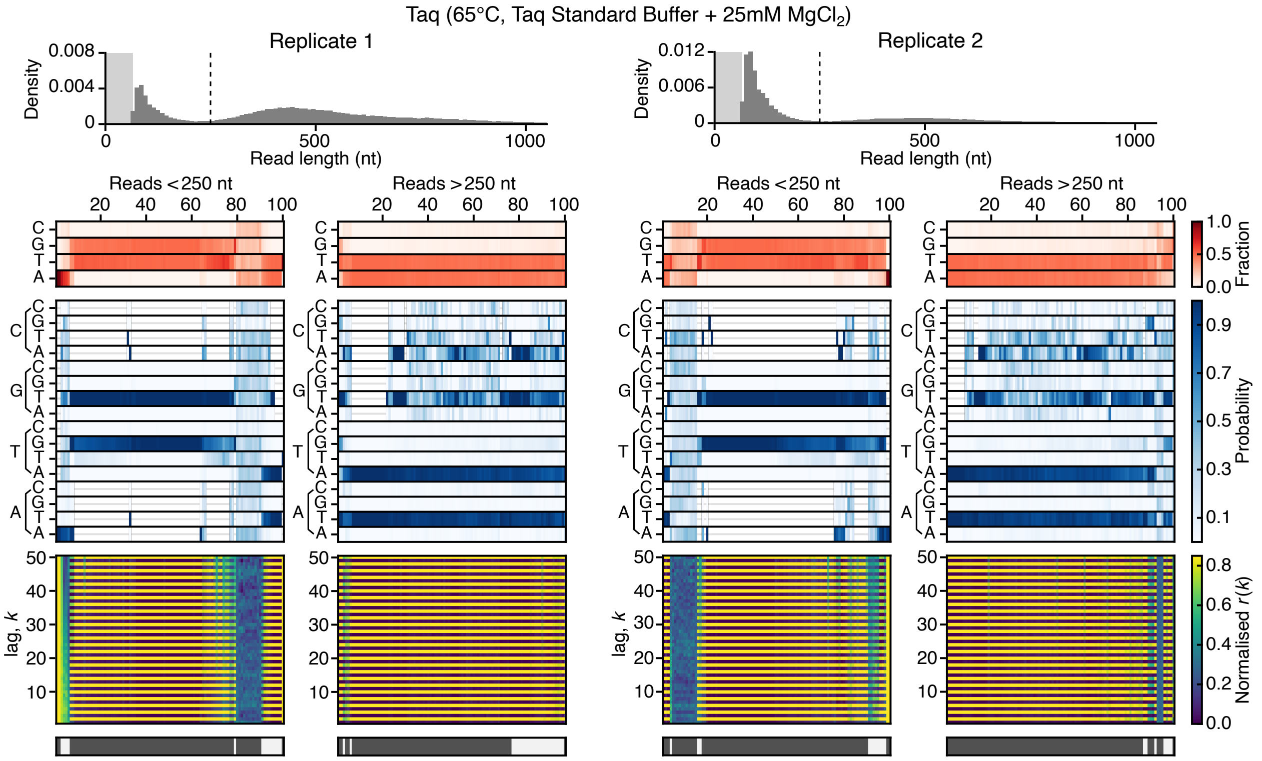


**Supplementary Figure 7: Sequence analysis of Taq (65°C, Taq Standard Buffer, 25 mM $MgCl_2$).** The top histograms show the sequence length distribution with the lightly shaded region denoting the 0–65 nt range and dashed line denoting the 250 nt read length. Below this, heatmaps show for a random subset of reads smaller and larger than 250 nt (left and right plots, respectively) the following information (top–bottom): 1. Sequence composition (red heatmap), 2. Probability of transitioning from one base to another (blue heatmap), 3. Autocorrelation analysis capturing the similarity of the sequence compared to itself after varying nucleotide shifts/lag *k* (blue to yellow heatmap), and 4. the seven top clusters of reads (alternating dark and light grey). Reads are displayed vertically and hierarchically clustered such that similar sequences are grouped.

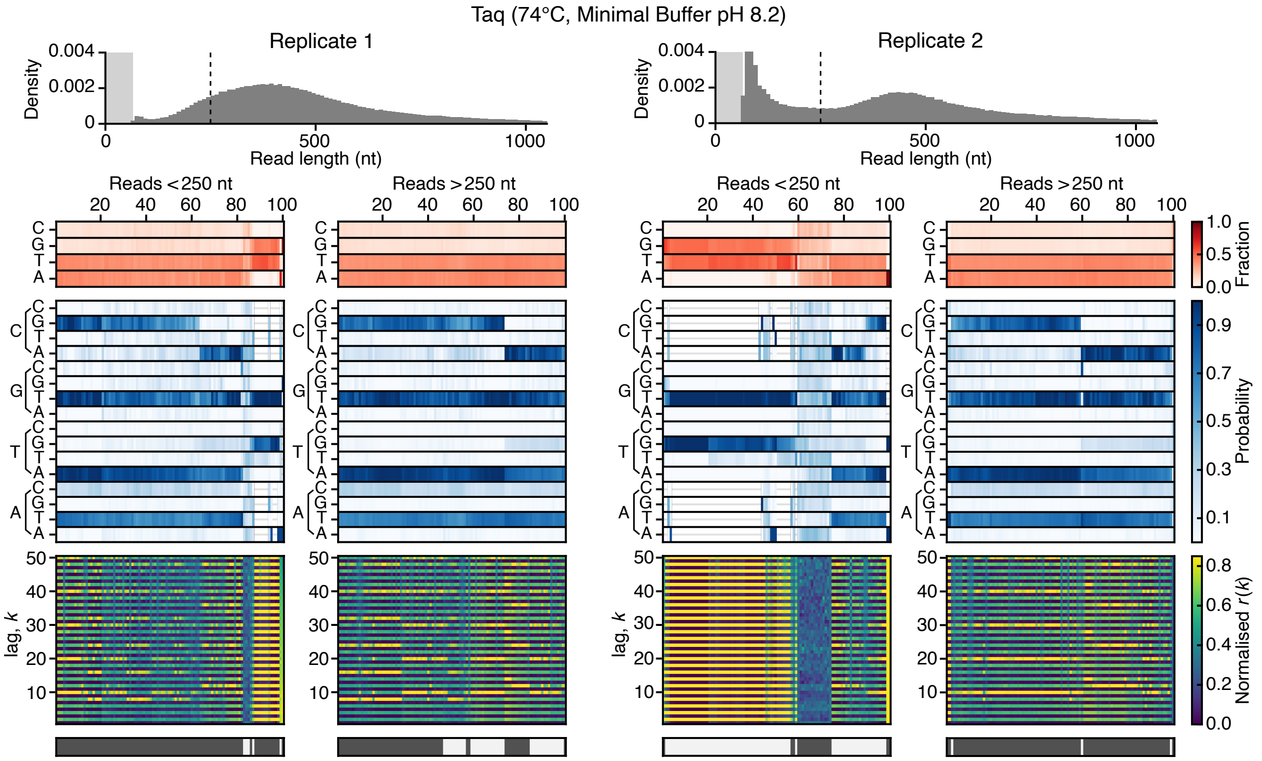


**Supplementary Figure 8: Sequence analysis of Taq (74°C, Minimal pH 8.2 Buffer).** The top histograms show the sequence length distribution with the lightly shaded region denoting the 0–65 nt range and dashed line denoting the 250 nt read length. Below this, heatmaps show for a random subset of reads smaller and larger than 250 nt (left and right plots, respectively) the following information (top–bottom): 1. Sequence composition (red heatmap), 2. Probability of transitioning from one base to another (blue heatmap), 3. Autocorrelation analysis capturing the similarity of the sequence compared to itself after varying nucleotide shifts/lag *k* (blue to yellow heatmap), and 4. the seven top clusters of reads (alternating dark and light grey). Reads are displayed vertically and hierarchically clustered such that similar sequences are grouped.

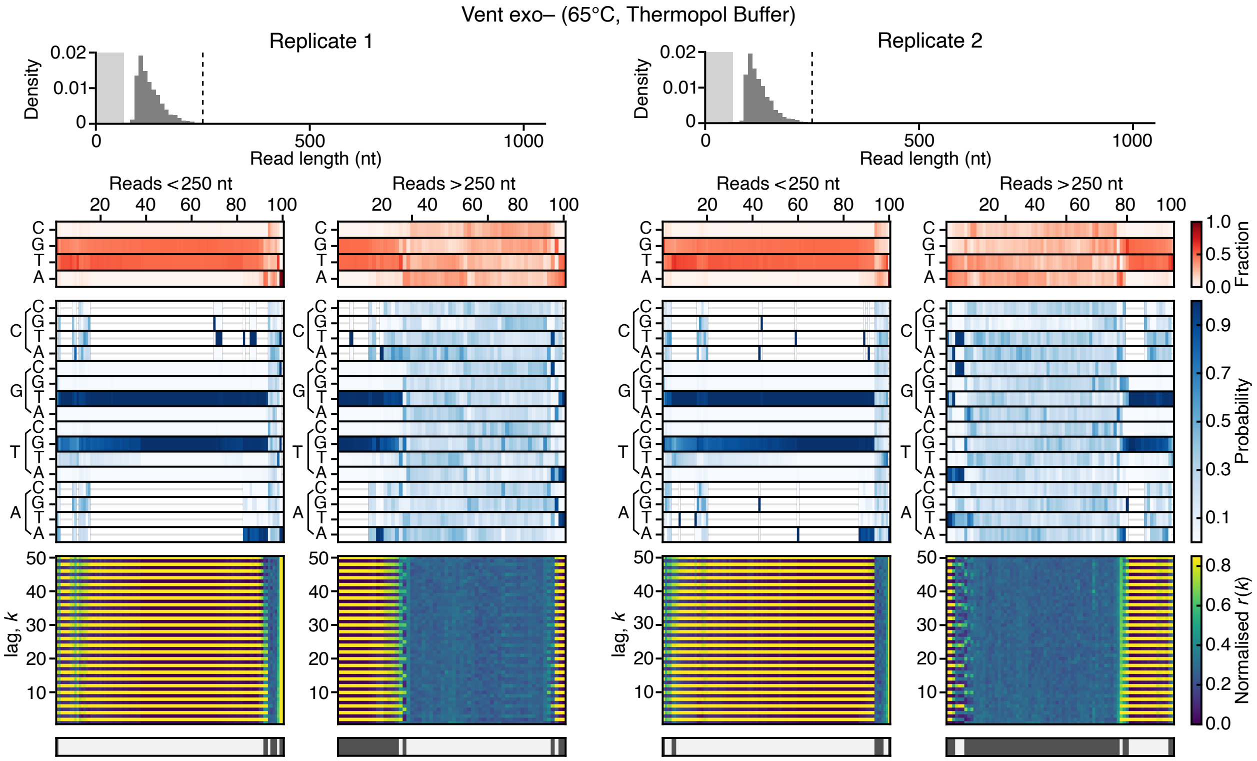


**Supplementary Figure 9: Sequence analysis of Vent exo– (65°C, Thermopol Buffer).** The top histograms show the sequence length distribution with the lightly shaded region denoting the 0–65 nt range and dashed line denoting the 250 nt read length. Below this, heatmaps show for a random subset of reads smaller and larger than 250 nt (left and right plots, respectively) the following information (top–bottom): 1. Sequence composition (red heatmap), 2. Probability of transitioning from one base to another (blue heatmap), 3. Autocorrelation analysis capturing the similarity of the sequence compared to itself after varying nucleotide shifts/lag $k$ (blue to yellow heatmap), and 4. the seven top clusters of reads (alternating dark and light grey). Reads are displayed vertically and hierarchically clustered such that similar sequences are grouped.

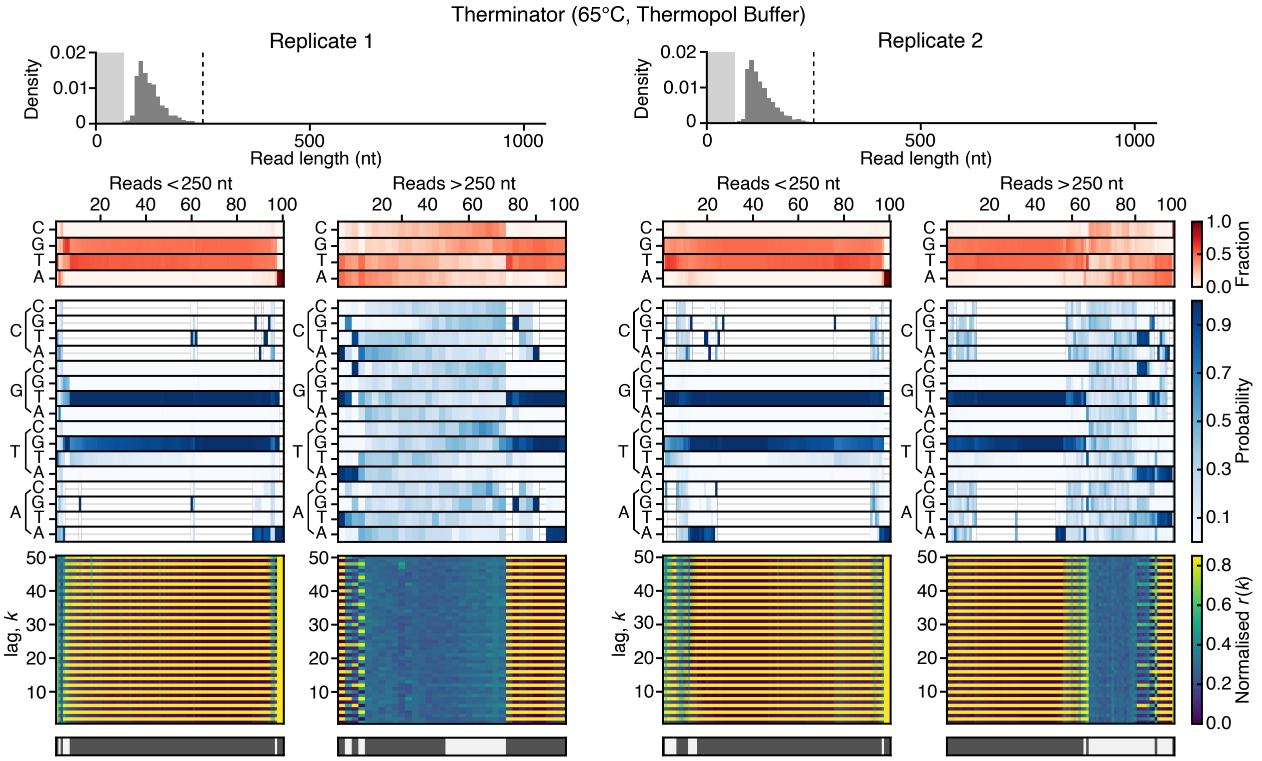


**Supplementary Figure 10: Sequence analysis of Therminator (65°C, Thermopol Buffer).** The top histograms show the sequence length distribution with the lightly shaded region denoting the 0–65 nt range and dashed line denoting the 250 nt read length. Below this, heatmaps show for a random subset of reads smaller and larger than 250 nt (left and right plots, respectively) the following information (top–bottom): 1. Sequence composition (red heatmap), 2. Probability of transitioning from one base to another (blue heatmap), 3. Autocorrelation analysis capturing the similarity of the sequence compared to itself after varying nucleotide shifts/lag *k* (blue to yellow heatmap), and 4. the seven top clusters of reads (alternating dark and light grey). Reads are displayed vertically and hierarchically clustered such that similar sequences are grouped.

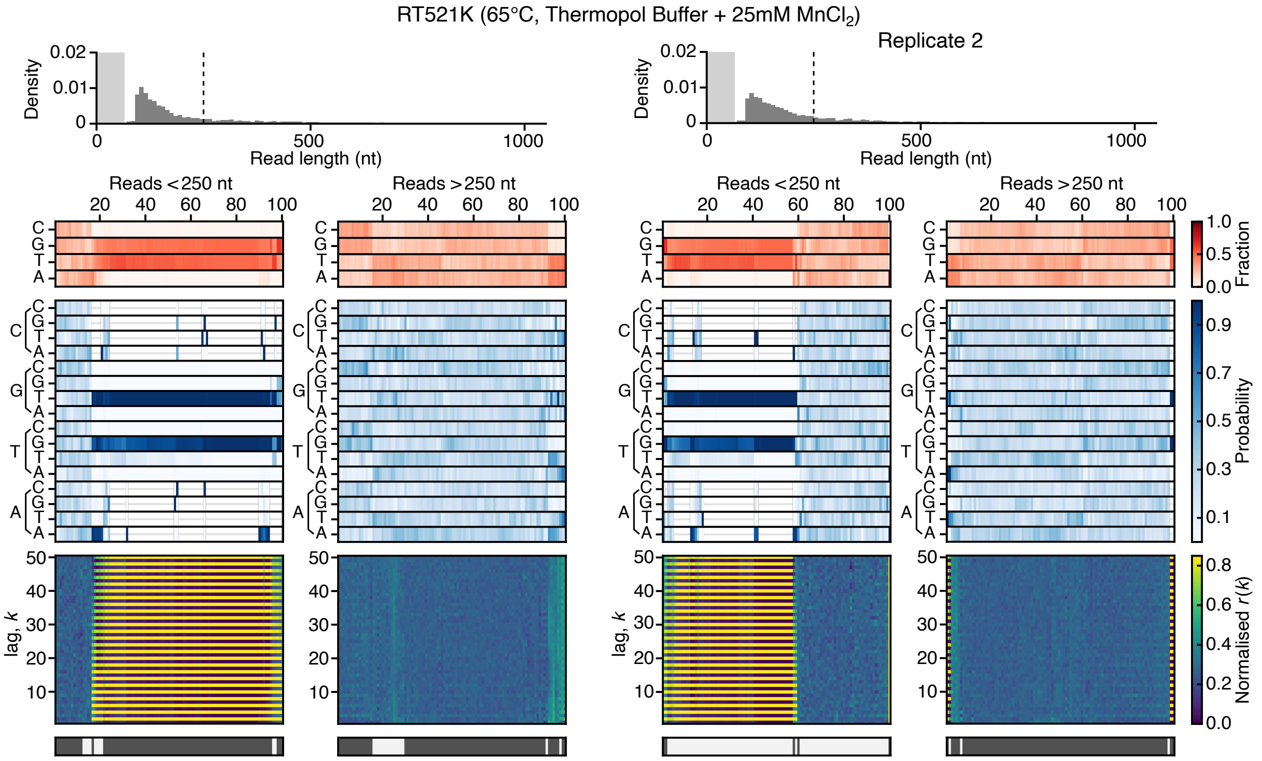


**Supplementary Figure 11: Sequence analysis of RT521K (65°C, Thermopol Buffer, 25 mM $MnCl_2$).** The top histograms show the sequence length distribution with the lightly shaded region denoting the 0–65 nt range and dashed line denoting the 250 nt read length. Below this, heatmaps show for a random subset of reads smaller and larger than 250 nt (left and right plots, respectively) the following information (top–bottom): 1. Sequence composition (red heatmap), 2. Probability of transitioning from one base to another (blue heatmap), 3. Autocorrelation analysis capturing the similarity of the sequence compared to itself after varying nucleotide shifts/lag *k* (blue to yellow heatmap), and 4. the seven top clusters of reads (alternating dark and light grey). Reads are displayed vertically and hierarchically clustered such that similar sequences are grouped.

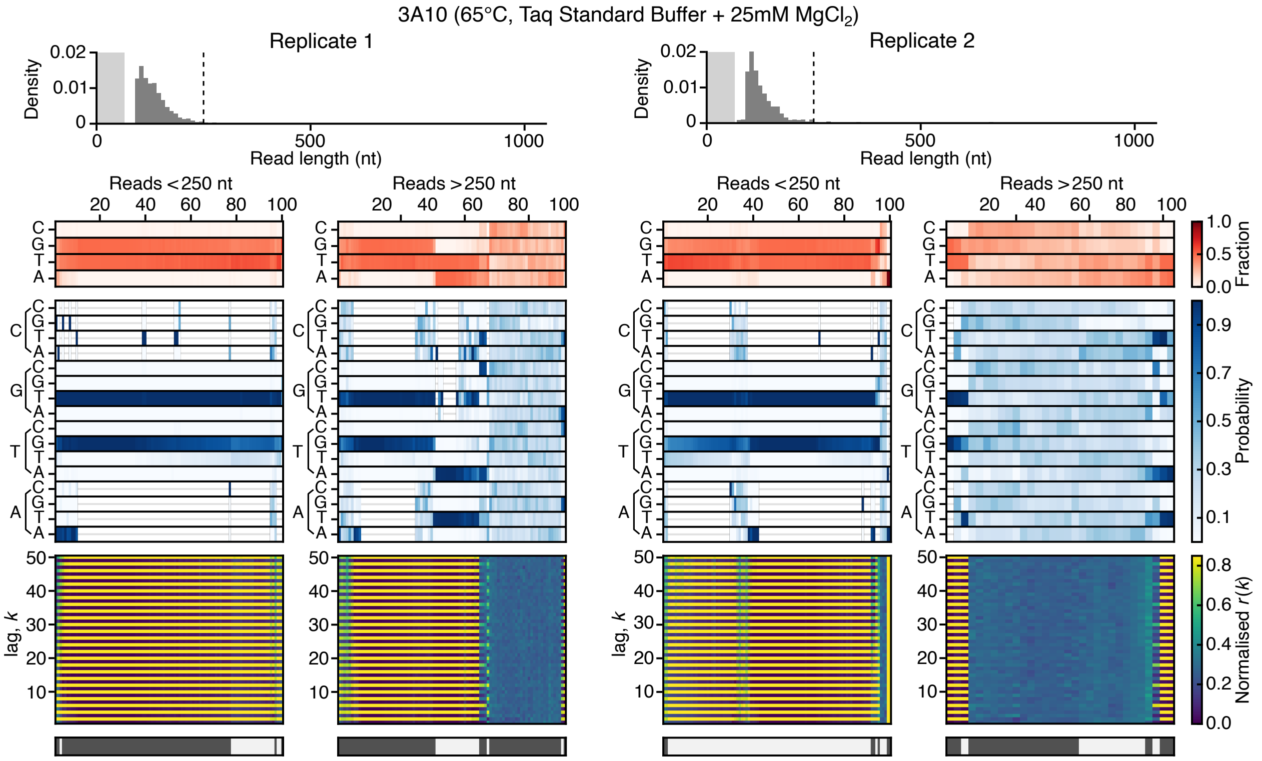


**Supplementary Figure 12: Sequence analysis of 3A10 (65°C, Taq Standard Buffer, 25 mM $MgCl_2$).** The top histograms show the sequence length distribution with the lightly shaded region denoting the 0–65 nt range and dashed line denoting the 250 nt read length. Below this, heatmaps show for a random subset of reads smaller and larger than 250 nt (left and right plots, respectively) the following information (top–bottom): 1. Sequence composition (red heatmap), 2. Probability of transitioning from one base to another (blue heatmap), 3. Autocorrelation analysis capturing the similarity of the sequence compared to itself after varying nucleotide shifts/lag $k$ (blue to yellow heatmap), and 4. the seven top clusters of reads (alternating dark and light grey). Reads are displayed vertically and hierarchically clustered such that similar sequences are grouped.

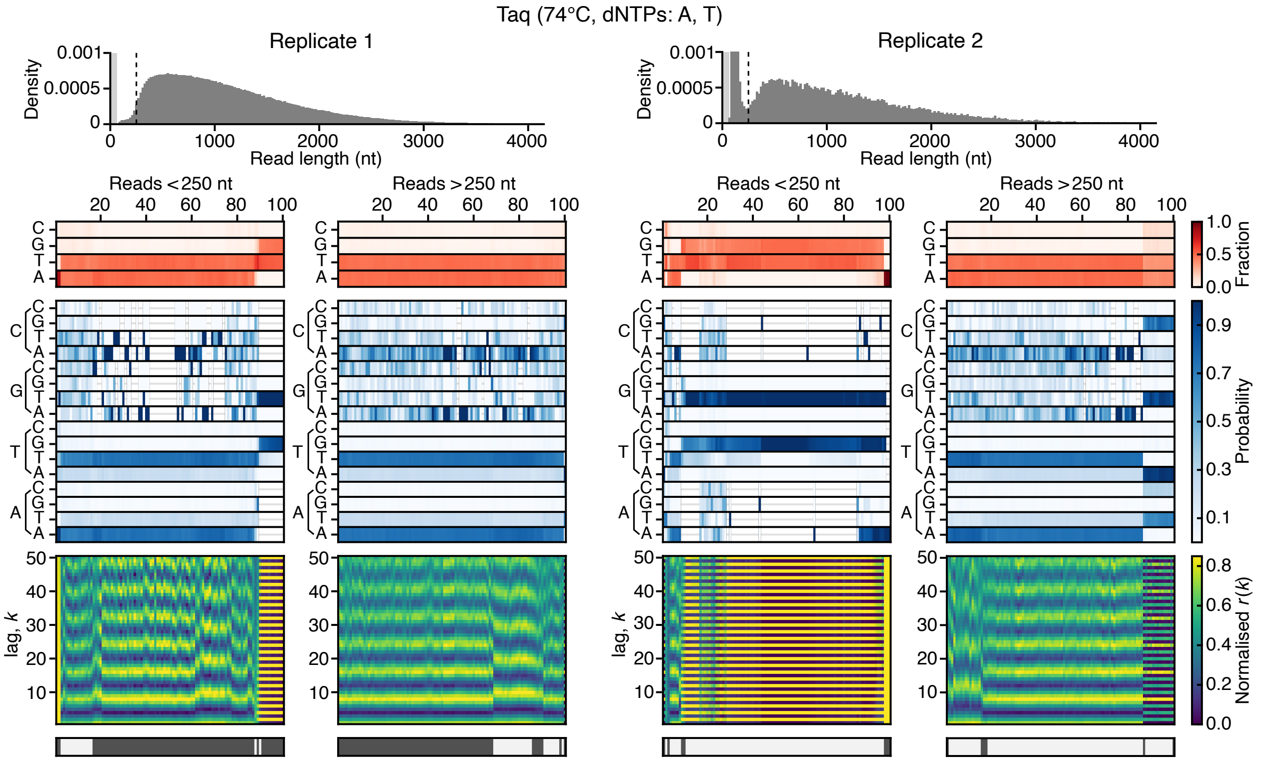


**Supplementary Figure 13: Sequence analysis of Taq (65°C, Taq Standard Buffer, only adenine and thymine provided).** The top histograms show the sequence length distribution with the lightly shaded region denoting the 0–65 nt range and dashed line denoting the 250 nt read length. Below this, heatmaps show for a random subset of reads smaller and larger than 250 nt (left and right plots, respectively) the following information (top–bottom): 1. Sequence composition (red heatmap), 2. Probability of transitioning from one base to another (blue heatmap), 3. Autocorrelation analysis capturing the similarity of the sequence compared to itself after varying nucleotide shifts/lag $k$ (blue to yellow heatmap), and 4. the seven top clusters of reads (alternating dark and light grey). Reads are displayed vertically and hierarchically clustered such that similar sequences are grouped.

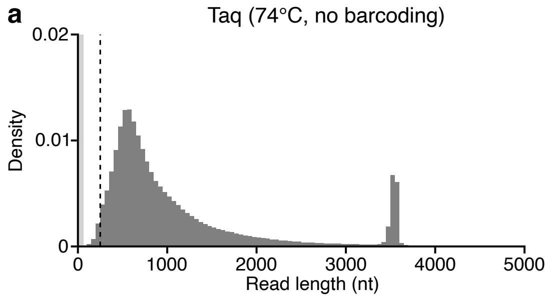


**Supplementary Figure 14: Read length distributions from non-barcoded sequencing runs.** (**a**) Taq at 74°C in Taq Standard Buffer. (**b**) Vent at 74°C in Taq Standard Buffer. histograms show the sequence length distribution with the lightly shaded region denoting the 0–65 nt range and dashed line denoting the 250 nt read length.

**Supplementary Table 1: Doodling activity of Taq with differing dNTP availability** [a]

| Polymerase | Temp. (°C) | Buffer | dNTPs | DNA mass [b] (ng/μL) | % Reads >250 nt | % Reads >1000 nt |
|---|---|---|---|---|---|---|
| Taq | 74 | Taq Standard | A | <4 | 48.6 | 11.8 |
| | | | | 15 | 16.6 | 3.9 |
| | | | T | <4 | 66.8 | 16.0 |
| | | | | 11 | 37.0 | 10.8 |
| | | | C | <4 | 45.5 | 10.8 |
| | | | | <4 | 27.7 | 7.2 |
| | | | G | <4 | 42.7 | 10.7 |
| | | | | <4 | 23.2 | 6.3 |
| | | | A, T | 23 | 98.4 | 50.3 |
| | | | | <4 | 80.9 | 42.2 |
| | | | A, C | 12 | 63.6 | 17.1 |
| | | | | <4 | 31.8 | 8.3 |
| | | | A, G | 7 | 80.3 | 19.9 |
| | | | | <4 | 48.3 | 12.5 |
| | | | T, C | 8 | 59.0 | 14.2 |
| | | | | <4 | 22.4 | 5.3 |
| | | | T, G | 21 | 48.2 | 11.3 |
| | | | | <4 | 34.5 | 9.1 |
| | | | C, G | 19 | 73.6 | 17.3 |
| | | | | <4 | 30.5 | 7.3 |
| | | | T, C, G | 16 | 60.1 | 15.3 |
| | | | | <4 | 20.7 | 4.2 |
| | | | A, T, C, G | 100 | 97.9 | 24.4 |
| | | | | 148 | 98.1 | 27.6 |

a. All reactions were 100 μL and run for 16 hours with DNA mass produced and read lengths given for both experimental replicates.

b. Values of <4 ng/μL were below the detection limit of the fluorometer used.
